# Large-scale cortical networks are organised in structured cycles

**DOI:** 10.1101/2023.07.25.550338

**Authors:** Mats W.J. van Es, Cameron Higgins, Chetan Gohil, Andrew J. Quinn, Diego Vidaurre, Mark W. Woolrich

## Abstract

The brain needs to perform a diverse set of cognitive functions essential for survival, but it is unknown how it is organized to ensure that each of these functions is fulfilled within a reasonable period. One way in which this requirement can be met is if each of these cognitive functions occur as part of a repeated cycle. Here, we studied the temporal evolution of canonical large-scale cortical networks, and show that while network dynamics are stochastic, the overall ordering of their activity forms a robust cyclical pattern. This cyclical structure groups states with similar function and spectral content at specific phases of the cycle and occurs at timescales of (300-1000 ms). These results are reproduced in five large magnetoencephalography (MEG) datasets. Moreover, we show that metrics that characterize the cycle strength and speed are heritable, and relate to age, cognition, and behavioural performance. These results reveal for the first time that the activations of a canonical set of large-scale cortical networks are organised in an inherently cyclical manner, ensuring periodic activation of essential cognitive functions.

## Introduction

The human brain fulfils many cognitive and homeostatic functions in a flexible and adaptive manner, which is essential for survival. Yet, it is unclear how it is organised to ensure that each of these are fulfilled within a certain time frame when the brain is in a non-structured temporal environment. One way in which this requirement can be met is if each of the cognitive functions occur as part of a repeating cycle. Since large-scale cortical networks, as studied through functional brain imaging, are thought to underlie specialised cognitive functions ^1–10^, we can examine the dynamics of these cortical networks to see if cyclical patterns exist.

Research into spontaneous brain activity recorded in wakeful rest using magneto- and electroencephalography (M/EEG)^1,11–13^ and functional MRI (fMRI)^14–16^ has shown that transitions between cortical networks in wakeful rest, or resting state networks, are non-random, and different levels of organisation have been observed. For example, multimodal evidence from MEG^11,17^ and fMRI^5,18^ suggests that the default mode network (DMN) and dorsal attention network (DAN), associated with an inward versus outward orientation of attention respectively, are anti-correlated and are unlikely to transition into each other directly. Moreover, recent results from fMRI show that the non-random transitions between resting state networks contain a hierarchical component, with clusters of brain states that are more likely to transition into each other within but not across clusters^14^. These asymmetries in transition probabilities between brain networks have further been shown to be more directional in states of higher awareness and in more physically and cognitively demanding tasks in both electrophysiology^21,23^ and fMRI. However, the existence of cyclical patterns between a full set of canonical large-scale cortical networks has not previously been shown.

Here, we investigated the temporal dynamics of large-scale cortical networks in multiple MEG datasets obtained during wakeful rest. We developed a new method for quantifying the transition asymmetries of these networks at a range of time scales, which showed that asymmetric transitions are ubiquitous in human brain activity. Moreover, while individual transitions were stochastic, together they produced a robust cyclical pattern of cortical network activations on 300-1000 ms timescales, an order of magnitude longer than the average lifetime of a single network. These patterns were reproduced in five independent datasets and robustly show a preferred position of each brain network in the cycle. Furthermore, we show that cyclical summary metrics are heritable, and relate to age, cognition, and behavioural performance. Together, these results are the first to show an overarching flow of cortical networks and suggest that cortical network activations are inherently cyclical, ensuring periodic activation of essential cognitive functions.

### Functional brain networks activate in structured cyclical patterns

To explore the temporal dynamics of large-scale functional brain networks in resting state MEG, we first conducted a secondary analysis of previously published results ^24^. This new analysis considered the longer-term patterns of resting state network (RSN) activity in the Nottingham MEG UK dataset (55 subjects) ^25^. Using hidden Markov modelling (HMM), the analysis (see methods) identified *K* = 12 states reflecting distinct brain networks with unique spatial configurations of power and coherence that reoccur at different points in time. States are inferred that best explain the multivariate distribution of activity across the entire brain whenever that state is active; states do not model any single spatial region in isolation, although spatially confined activity may nonetheless be characteristic of a particular state. The state descriptions of all network states are shown in Figure S1.

States are inferred that best explain the multivariate distribution of activity across the entire brain whenever that state is active; states do not model any single spatial region in isolation, although spatially confined activity may nonetheless be characteristic of a particular state.

We characterised the tendency of network states to follow or precede each other using a novel method called Temporal Interval Network Density Analysis (TINDA; Figure 1). This method focusses on the variable-length intervals between network state occurrences, which relaxes more common assumptions of fixed-length timing patterns, an approach that we show is crucial to its success (see Figure S4). For each reference state, TINDA takes all intervals between reoccurrences of the same state (i.e., *state- -to-intervals)* and partitions them evenly in half. It then defines the Fractional Occupancy (FO) Asymmetry as the difference between the probability of another network state occurring in the first half versus the second half of those intervals. This measure captures if there is a tendency for a network state to follow, or precede, another state over variable timescales (Methods and Figure 1D-E).

**Figure 1.**
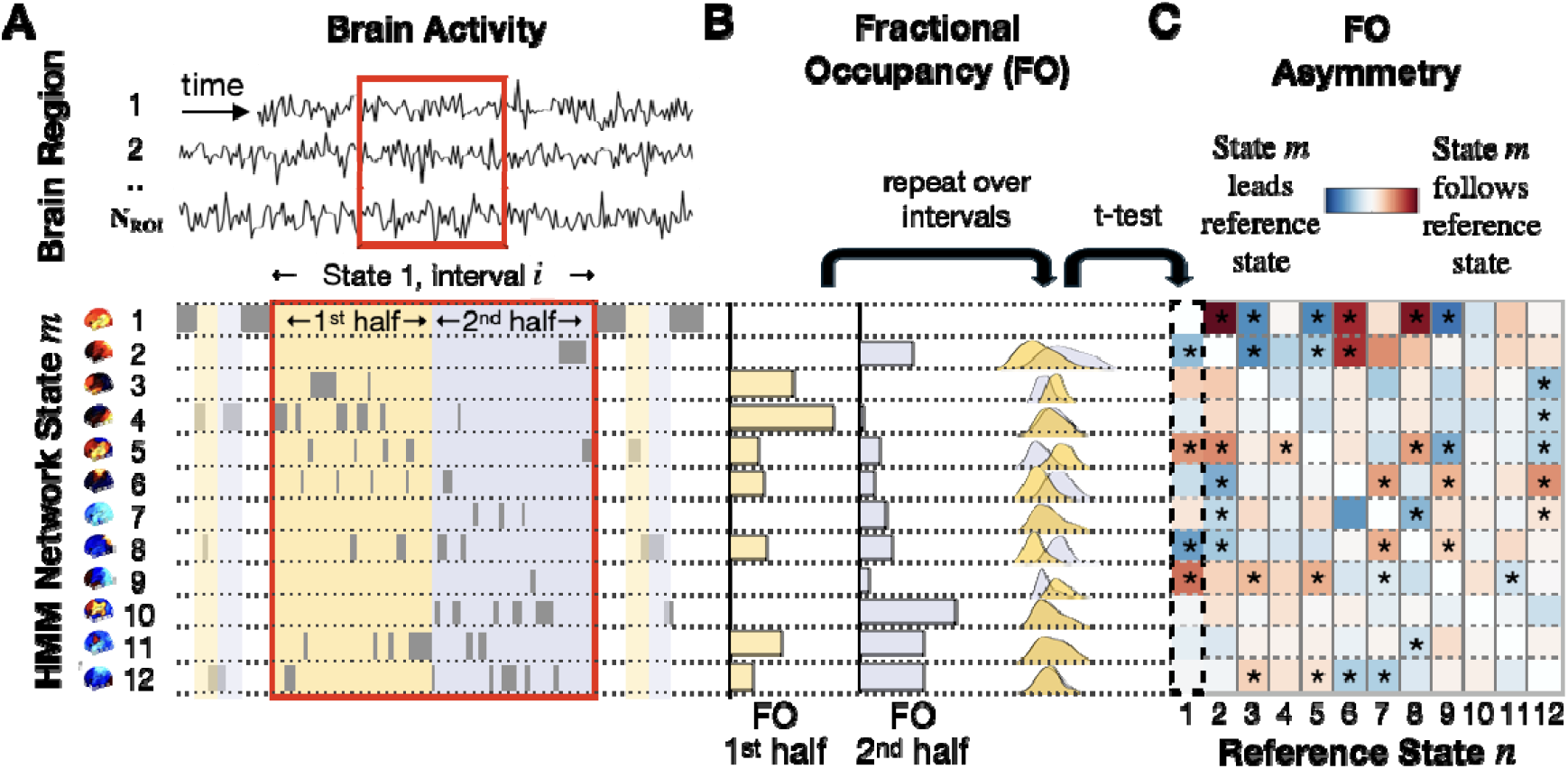
Schematic of Temporal Interval Network Density Analysis (TINDA*), with state* 1 as an example reference state; note that each state gets used in turn as the reference state, with the outputs then combined within the TINDA procedure. A) A segment of the (1 to N_ROI_) multi-region, resting state MEG data (top), and the inferred HMM state activations (bottom). Segment ***i*** is the period between reactivations of network state 1, which is further subdivided into two interval halves (1^st^ half: yellow; 2^nd^ half: blue). B) The fractional occupancy (FO, i.e., relative time spent) for each network state in both intervals in A (left), and the FO distributions over all state 1-to-state 1 intervals (right). C) The FO Asymmetry matrix shows the mean FO difference over intervals between the two interval halves with respect to a reference state (in this example state 1). This procedure is repeated for all reference states ***n*** to create the full FO asymmetry matrix, which is used in the results going forward. Asterisks denote significant (Bonferroni-corrected) FO asymmetries.

We used this method to investigate whether an overarching pattern emerged when every state’s tendency to follow or precede every other state was analysed. To illustrate its use more clearly, we first used this method on the intervals defined by subsequent visits to state (Figure 1). This revealed that certain network states (states 5, t(54) = 5.1, p=4.1*10^-6^; and 9, t(54) = 6.4, p=3.7*10^-8^) tend to occur after the state 1, while other states (states 2, t(54) = -4.6, p=2.3*10^-5^; and 8, t(54) = -6.1, p=9.9*10^-8^) tend to occur before the state 1. All other states (3, 6, 7, 10, 11, and 12) did not exhibit significant asymmetric activation probability after Bonferroni correction for multiple comparisons. In the interest of reproducibility, we repeated the same analysis for the equivalent state in two other large datasets (Cam-CAN (N=612) ^26,27^ and HCP (N=79) ^28^, and found consistent results (see Figure S2).

We next investigated whether asymmetries in activation probabilities also exist for other network states. Using TINDA on all pairs of network states, we confirmed that this was indeed the case, and moreover, that these pathways were unique to each state (Figure 1C, and Figure S2). All results that follow rely on the full FO asymmetry matrix (Figure 1C), i.e., where TINDA is applied to state-*n-to-n* intervals for *n* ∈ 1: *K*.

We then explored the possibility that the asymmetries in network activation probabilities are unified by an overarching structure. In particular, visual inspection of these networks raised the possibility they were unified by a globally cyclical structure (Figure 2D), an emergent dynamic that could not arise trivially from the first-order state asymmetries (p<0.01, see Supplement II). We defined the cycle strength (***S*)** to test the potential cyclical structure statistically (see methods for details). Cycle strength is +1 for graphs where all transitions are perfectly clockwise, zero for completely stochastic graphs, and negative for overall counterclockwise transitions (note that when states are ordered to maximise ***S***, negative cycle strength can only be true for individual subjects, not for the group average; and vice versa when ***S*** is minimised). We confirmed that the cyclical pattern as a result of all FO asymmetries together could not have arisen by chance by permuting network state labels within each subject. In each of 1000 permutations, the order of states was shuffled independently for each subject and cycle strength was computed using the optimal cycle order for that permutation); the observed cycle strength was significantly greater than in permutations (mean (standard deviation): ***S*** = 0.066 (0.041); p<0.001). Moreover, in additional control analyses, we ruled out the possibility that the cyclical pattern could arise from common (rhythmic) physiological artefacts, see Supplement VI.

**Figure 2.**
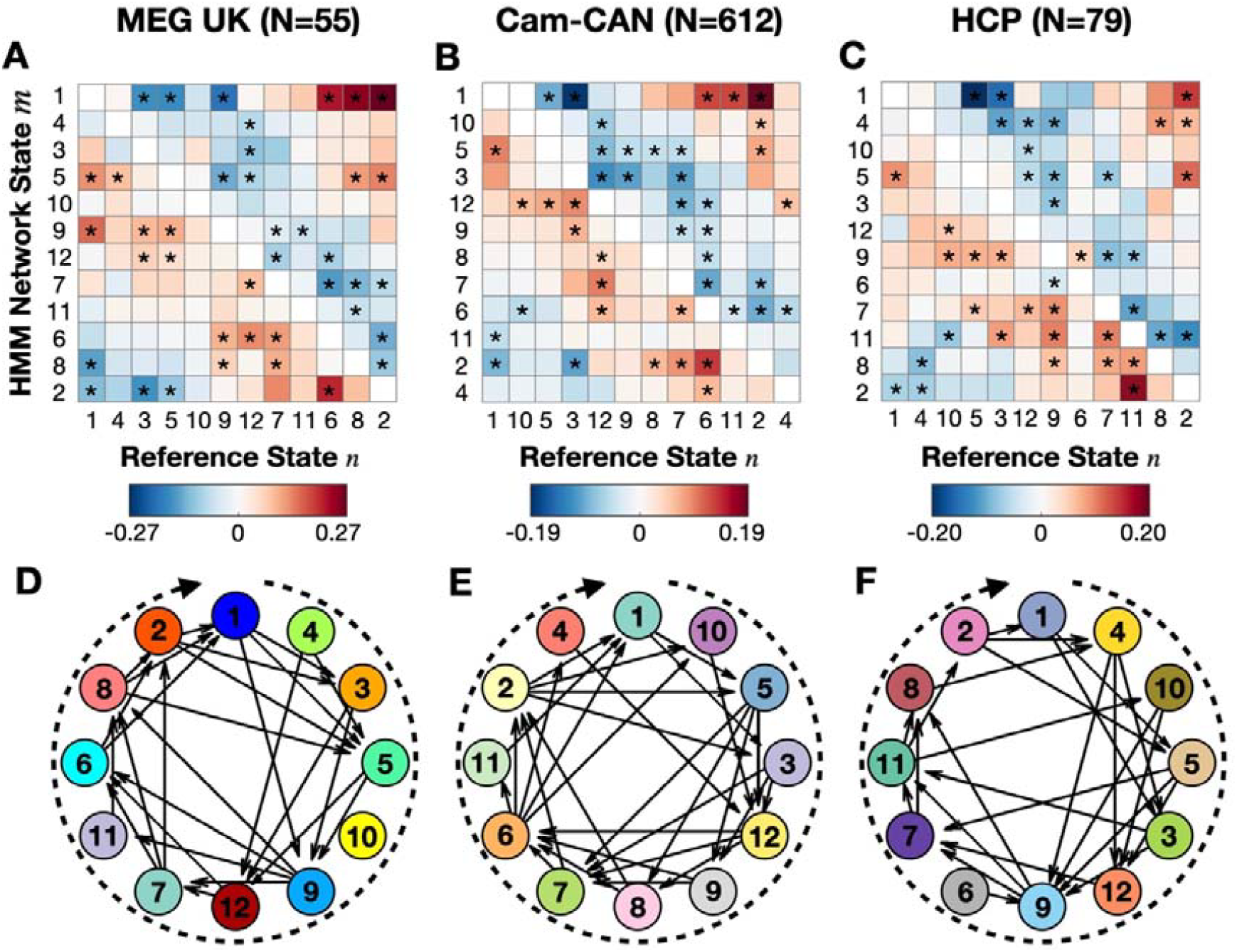
Reordering the states to optimise the flow of pairwise FO asymmetries reveals an overarching cyclical activation structure of functional brain networks in three large MEG datasets (MEG UK: left, Cam-CAN: middle, HCP: right) A-C) The group-average Fractional Occupancy (FO) Asymmetry describes the activation probability of one network state (y-axis) relative to another (x-axis). Asterisks denote statistically significant elements, shown in D-F) as edges in a directed graph. The colours of nodes in the different datasets are distinct to indicate that network state descriptions are inferred independently from each dataset. State numbers in Cam-CAN/HCP are matched to the MEG UK dataset (see Methods), which is numbered in order of decreasing coherence.

In the interests of reproducibility, we replicated these analyses in the two other datasets, confirming both the presence of cyclical dynamics and the consistency of individual state ordering within the cyclical configuration across all datasets. HMMs were independently trained on each dataset, after which states were reordered to match the ordering in the MEG UK dataset (see Methods); state numbers across the three datasets thus refer to equivalent network states. We confirmed that cycle strength was higher than permutations in both Cam-CAN (***S*** = 0.049 (0.033); p<0.001; Figure 2B/E), and HCP (***S*** = 0.048 (0.035); p<0.001; Figure 2C/F). This confirmed the presence of a cyclical structure in all three datasets, but it remained plausible that these were *different* cyclical structures. Because we identified equivalent states in all three datasets, we could test whether the order of states in the cycle was the comparable between datasets. We computed the cycle phase difference between equivalent states in each dataset and compared this with a random placement of states across the cycle (i.e., permuting state positions 10000 times). This analysis confirmed that the order of states in the Cam-CAN cycle matched the order in MEG UK: the mean phase difference (Δθ) between equivalent states was smaller than expected by chance (Δθ = 0.645 rad, p = 0.0038; also see Supplement III). Despite the use of an entirely different parcellation in HCP, the same was true in this dataset (Δθ = 0.472 rad, p = 0.0001). These analyses thus show that the *same* cyclical dynamics can be observed across three independent datasets.

### Cyclical structure is strongest over timescales of seconds

Given the strength of this cyclical activation pattern, we considered why it had not been characterised previously in the literature. TINDA differs from other methods of characterising dynamics in that it measures dynamics over inter-state intervals of variable length. These intervals have a highly dispersive distribution with a very long tail ^11,24,29^. Common means of modelling temporal dynamics typically assume either a Markovian structure, as in our work, (i.e., that the state at one timepoint is conditionally dependent on only the immediately prior state^11^) or a structure of temporal dependency with fixed length time-lags^30^. Simulations from either of these models trained on the existing dataset illustrates why such a cyclical activation pattern would not have been detected in previous work without an additional post-hoc analysis such as TINDA to capture dependencies that are not reflected explicitly in the model parameters. These simulations capture only a small part of this inherent cyclical structure, most of which is lost due to the variability of ISI durations (Figure S4). The fact that other models capture only a small part of this inherent cyclical structure underlines the significance of our novel approach.

This also suggests two key temporal features of the cyclical patterns we have characterised; firstly, that these cyclical patterns are instantiated over longer time scales; and secondly that they do not have a regular cyclical period (see Supplement VI). To verify this quantitatively, we looked at the dependency of the FO asymmetry and cycle strength on the interval duration (i.e., with in figure 1, the interval time between subsequent visits to the network state of interest ()). We expected that if cyclical patterns are instantiated over longer timescales, then the FO asymmetries and the characteristic cycle would only be apparent at longer interval times.

To do this we partitioned the distribution of interval times (Figure 3A) into five equally sized bins. We did this separately for each state to ensure there was no state bias in each bin. This procedure resulted in each bin containing an average (standard deviation) of 885 (111) intervals for each subject. We then reran the TINDA procedure separately on (the intervals from) each bin. Figure 3B shows that group mean cycle strength is close to zero for the bins with the shortest duration intervals and increases for bins with higher interval durations. Cycle strength is significantly higher than in permutations (see Methods) in bins 2-5, and strongest in the bin containing the longest interval durations (with a mean interval time of 3 seconds). This was replicated in the Cam-CAN and HCP datasets (Fig S5), and together these results prove that the cyclic activation of network states is occurring at timescales on the order of seconds.

**Figure 3.**
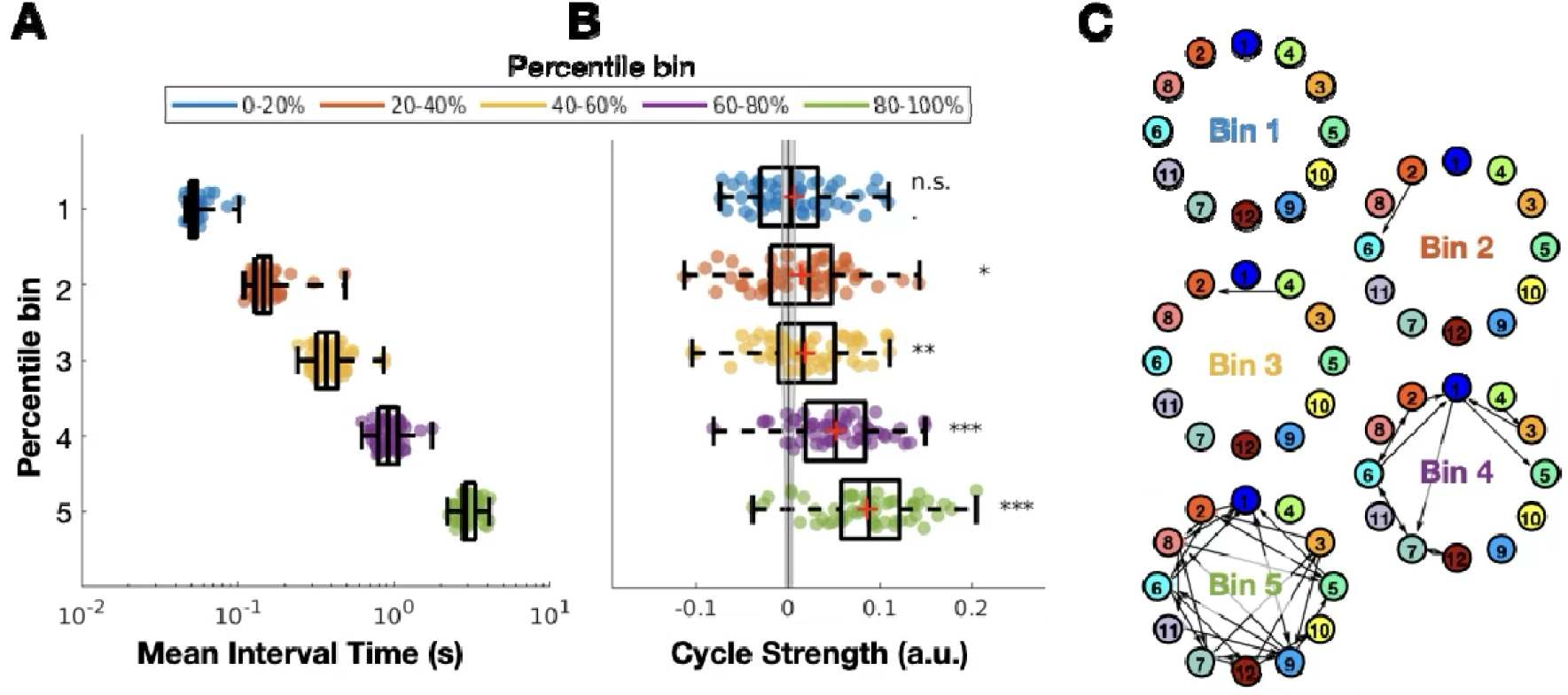
The observed cyclical organisation of network state activations is driven by longer interval times in the MEG UK dataset. For each subject and state, intervals were binned by interval times into five percentile bins. A) The mean duration of interval times over all states within each percentile bin. B) The cycle strength resulting from running TINDA on each percentile bin in A. Circles are individual subjects; boxplots display the median, mean (+), 25^th^ and 75^th^ percentile, and whiskers indicate the minimal and maximal value. The line and error bar around zero cycle strength are the mean and standard deviation of the empirical permutation distribution. Significant cycle strength is denoted by asterisks: * p<0.05, ** p<0.01, *** p<0.001, and “n.s.” denotes not significant. C) Graphs similar to Figure 2D for binned interval times with increasing duration from top to bottom. Note that arrows are only shown for significant FO asymmetries, i.e., Bin 1 does not contain any significant asymmetries and is therefore empty.

### Cyclical structure groups networks with similar spectral properties and function

Having established that resting state networks tend to activate in a cyclical progression, we next characterised what a complete traversal of a single cycle might look like. We did this by mapping the spatial/spectral network state descriptions provided by the HMM onto the cycle. The result of this is shown in Figures 4 for the MEG UK dataset with power maps, and S7 for the other datasets and coherence maps. We emphasise that each network state comprises a spatially defined pattern of power and coherence. To display these high dimensional representations more succinctly, Figure 4A only shows the single dominant spatio-spectral mode in each state (see Methods and Fig S1); this information is further condensed and summarised in figures 4B-C. Quantitative comparisons of the MEG HMM states and the Yeo7 atlas^31^ have been made in Supplement XII.

**Figure 4.**
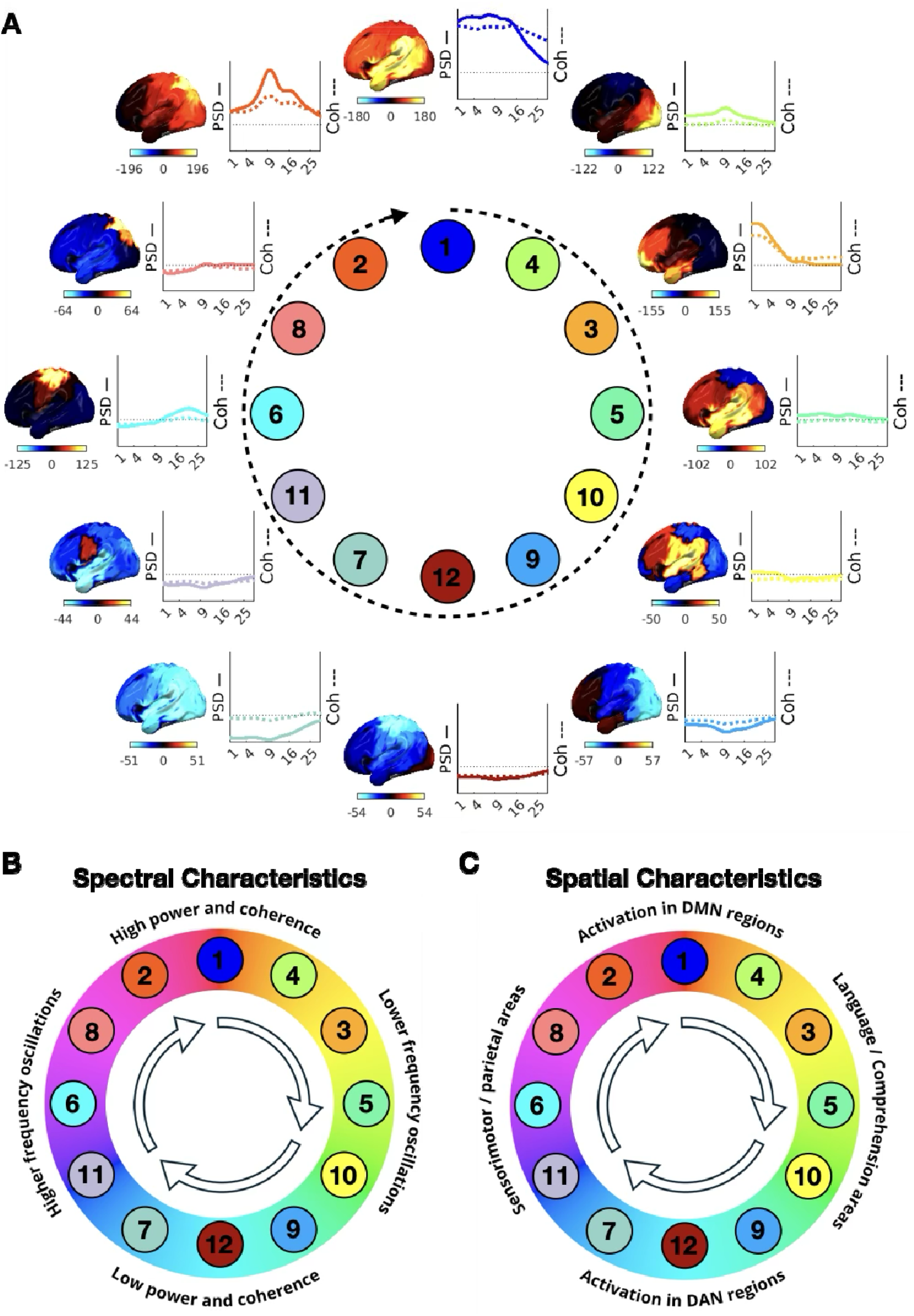
The cyclical structure groups together network states that have similar spectral properties and cognitive function in the MEG UK dataset. A) The spatio-spectral characteristics of functional brain networks are embedded in their cyclical progression (see main text). Each brain map shows the percentage increase in power (for visualisation purposes, shown relative to the mean over states), projected onto the left hemisphere (see Fig. S7 for the coherence maps and replication in Cam-CAN and HCP, and Fig S1. for detailed spectral characteristics for each network state). To the right of each brain map is the spatial average PSD (solid line) and coherence (dotted line) as a function of frequency, relative to the average over states (horizontal dotted red line). B-C) Qualitatively summarising the spectral (B) and spatial (C) modes seen in A (also see Supplement XII) .

The first major mode of differentiation between network states emerges on the North-South axis of the clock face. States in the upper quadrant have a higher overall power and inter-area coherence (i.e., phase locking). States 1 and 3 in particular show strong overlap with areas overlapping the DMN (including bilateral inferior parietal lobe, medial prefrontal cortex, and medial temporal lobe, also see Supplement XII). This is not a mere broadband power increase, but rather reflects different combinations of oscillatory activity in distinct or overlapping frequency bands ^29^. On the lower quadrant, states have lower overall power and inter-area coherence, particularly in sensorimotor and parietal areas. These states are associated with sensorimotor processing (state 9, 12) and the dorsal attention network (DAN), see Supplement XII.

A second mode of differentiation emerges on the East-West axis of the clock face. In terms of spectral activity, network states on the left of the quadrant display activity in higher frequency bands; for instance, state 6 is associated with beta band (14-30 Hz) activity, and state 2 with alpha band (7-13 Hz) activity. On the other hand, states on the right-hand side show activity in lower frequency bands, particularly the delta (1-4 Hz) and theta (4-7 Hz) band. Spatially, states on the left-hand side show increased low frequency activity in sensorimotor and parietal areas, which are associated with sensorimotor inhibition. Meanwhile, states on the right-hand side show activity mostly in fronto-temporal and language areas ^2,29,32^.

The differentiation in spatio-spectral activity suggests different types of brain function and processes are localised to particular phases of the cycle. For example, these results suggest that network states going into the DMN are linked to sensorimotor inhibition through increased alpha/beta power. In contrast, networks going away from the DMN comprise of slower frequency content in higher order fronto-temporal areas, which is followed in turn by low power sensorimotor states, and, in particular state 7, characterised by a decrease in oscillatory power in the parietal regions overlapping the DAN.

In the interest of reproducibility, this plot has been replicated on the Cam-CAN and HCP datasets. The main findings summarised in figure 4B and 4C were reliably reproduced (Figure S7), despite some moderate differences in network state definitions.

### Cycle statistics relate to cognition and demographics

Inspired by this qualitative segmentation of cycles into four “meta states” of distinct spatio-spectral characteristics, we defined a full cycle traversal as the sequential activation of these (See Methods and Supplement VIII for details). This allowed us to define cycle duration as a metric to summarise the time scale of these dynamics. Cycle duration was on average on the time scale of 300-1000 ms (MEG UK mean (µ) ± standard deviation (SD) = 549 (154) ms; Supplement VIII, Cam-CAN µ (SD) = 355 (62.4) ms; Figure 5D, HCP µ (SD) = 528 (104) ms). We could then relate cycle duration, or in fact its more normally distributed inverse (i.e., cycle rate), to individual traits, together with the previously defined cycle strength.

**Figure 5.**
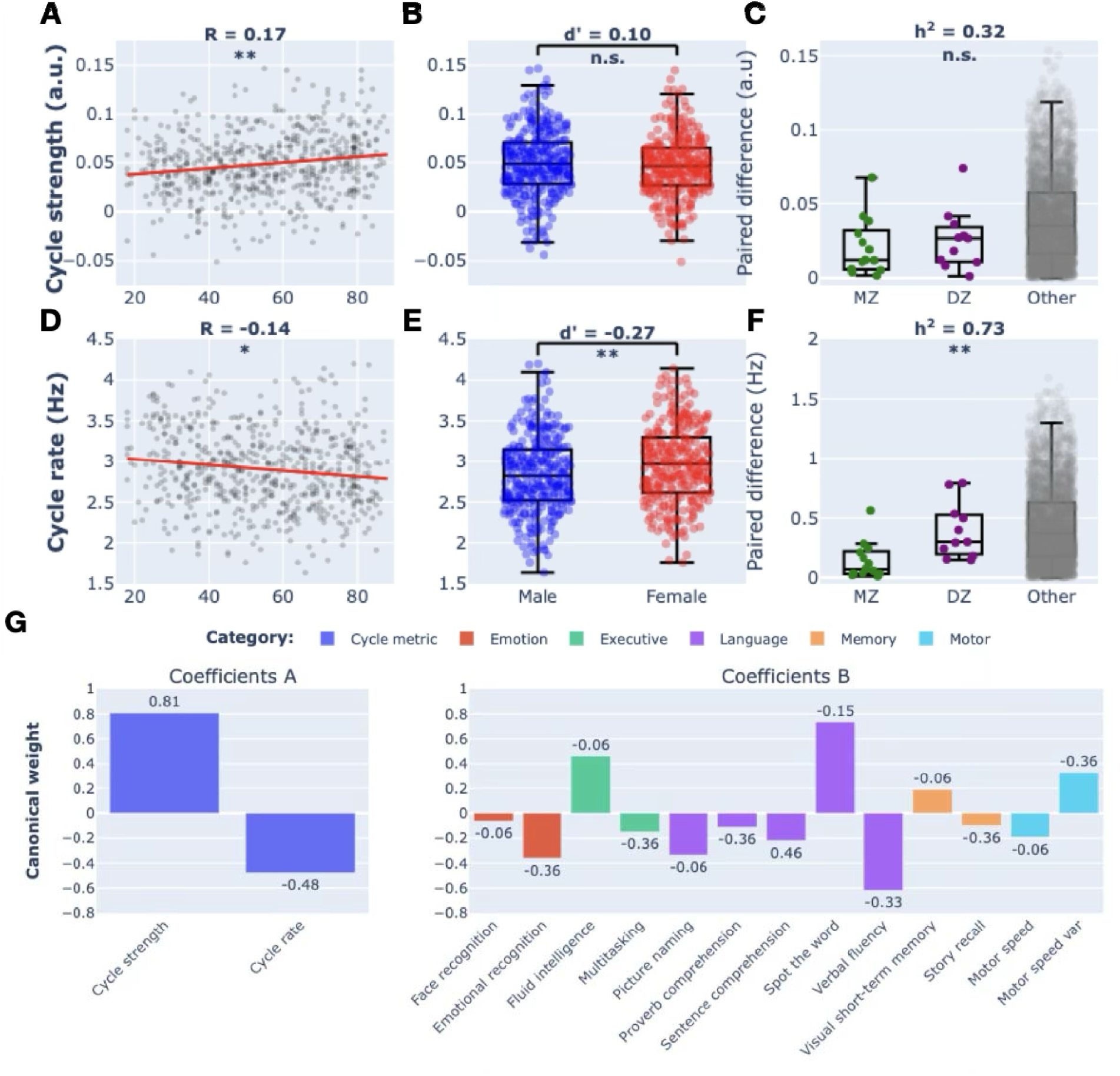
Cyclical activation statistics relate to individual traits and are heritable. A) Cycle strength as a function of age. The histogram on the right shows the distribution of cycle strength, and the group mean in red. B) Cycle strength as a function of sex. C) The absolute difference in cycle strength between monozygotic (MZ; “identical”) twins, Dizygotic (DZ; “non-identical”) twins or unrelated subject pairs. Circles correspond to individual subjects (A/B/D/E), or pairs of subjects (C/F); boxplots display the median, 25^th^ and 75^th^ percentile, and whiskers indicate the minimal and maximal value not considered outliers. n.s. denotes not significant, * p<0.05, ** p<0.01, *** p<0.001. D-F) As in A-C but for cycle rate (i.e., the inverse of cycle duration). G) Canonical weights from the second (significant) pair of canonical variates; cycle metrics (left) and cognitive scores (right). See Supplement IX for evidence these results are robust to minor changes in metric definitions, and Supplement XIII for the other canonical variate variables and post-hoc Pearson correlations between individual variables.

We first made sure that these cycle strength and cycle rate are consistent within individuals. We computed the intraclass correlation coefficient (ICC) on the metrics for the three sessions per subject available in the HCP dataset. This confirmed that both metrics are consistent across sessions, (cycle strength: r = 0.43 (95% CI: 0.29-0.56), F(78,158) = 3.2, p = 1.9 * 10^-10^; and cycle rate: r = 0.80 (95% CI: 0.72-0.86), F(78,158) = 12.9, p = 0). We also found that these metrics are robust to the number of network states fitted in the first level HMM (Supplement IX). We then took advantage of the large and equally distributed age range (18-86 years) and sex in the Cam-CAN dataset and asked whether either could be predicted by cycle strength or cycle rate (Figure 5). Because both age and sex are known to affect heart rate, and the heartbeat has a strong effect on the MEG signal, we first regressed out heart rate. Next, we fitted a General Linear Model (GLM) which revealed that cycle strength reliably predicted age (beta = 2.49, SE = 0.75, t(605) = 3.30, p = 0.0010; post-hoc Pearson correlation R = 0.16), but not sex (beta = -0.052, SE = 0.085, t(605) = -0.61, p = 0.54; Cohen’s d = -0.10), and cycle rate predicted age (beta = -2.04, SE = 0.75, t(605) = -2.71, p = 0.0070; post-hoc Pearson correlation R = -0.15), and sex (beta = 0.24, SE = 0.086, t(605) = 2.81, p = 0.0050; Cohen’s d = 0.27). These findings are robust to minor changes in the way that metrics are defined (see Supplement IX). We replicated these correlations in the other datasets and confirmed the correlations between age and cycle rate and strength but found no statistical difference between males and females (Supplement X). A post-hoc analysis revealed that the correlation between age and cycle strength can be explained by a combination of 1) stronger pairwise asymmetries between network states on average, and 2) fewer deviations from the cycle structure (i.e., fewer backwards or random transitions) (Supplemental Figure 11).

The correlations between cycle metrics and age suggest that older people have slower, and stronger cycle dynamics. Given cognitive slowing and inflexibility is often observed in older people ^14,33^, we wondered whether these were related. We first regressed out age, sex, and heart rate from all variables, and then used a canonical correlation analysis (CCA) to find a relationship between cycle metrics and cognitive scores, resulting in two, orthogonal canonical correlation variates. This confirmed a statistically significant relationship between cognitive scores and cycle metrics for the second (R = 0.17, F(12, 597) = 1.51, p = 0.0087 versus permutations; Figure 5G), but not the first (R = 0.19, F(26, 1192) = 1.54, p = 0.19; Figure S15) variate. Notably, the canonical weights of cycle metrics for the significant relationship with cognitive scores were in the opposite direction, and so was the correlation between these metrics and age. This could suggest a relationship between cycle dynamics and age-related cognitive decline. Replication of this finding was not assessed in other datasets because comparable cognitive scores were not available.

We next wondered whether cycle metrics could be genetically determined. The Cam-CAN dataset did not allow us to test this for the lack of twin data, so we turned to the HCP dataset, which contains data of mono- and dizygotic twins, and unrelated pairs of subjects using an ACE model of heritability ^34,35^. The ACE aims to partition the phenotypic variance into three components: additive genetic variance (A), shared environmental factors (C), and nonshared environmental factors (E). Despite the relatively small cohort of twin data, we found strong evidence that cycle rate, but not cycle strength, is heritable (Figure 5C/F). In fact, 73% of the variance in cycle rate in the population could be explained by genetic factors (h^2^ = 0.73, 95% CI = 0.29-0.98, p = 0.0039). We did not find such an effect for cycle strength (h^2^ = 0.32, 95% CI = 0.01-0.67, p = 0.12), nor did we find evidence that environmental factors affected cycle metrics (cycle strength: c^2^ = 0.18, 95% CI = 0-0.41; cycle rate: c^2^ = 0, 95% CI = 0-0.43). To make sure demographic or morphometrics factors did not bias these results, we systematically regressed out potentially confounds (e.g., age, sex, brain volume, etc.; see Methods and Figure S10). The heritability estimate of cycle rate remained high (h^2^ = 0.68) and significant even with the most stringent confound modelling.

### Preserved cycle structure in task data is behaviourally relevant

Having established that cortical networks activate in cycles across multiple datasets in a manner predictive of individual traits, it remained possible that they nonetheless reflect some neurophysiological feature of little or no relevance to cognitive processes. We therefore first asked whether the cyclical patterns observed during rest were related to spontaneous memory replay. Secondly, we tested whether cyclical patterns persisted in task data. and whether variance in cyclical metrics over task epochs related to variance in task performance.

In the memory replay ^36^ dataset, participants learned sequence structures between different visual images. The representations of these have been shown to replay spontaneously during a subsequent rest period ^36^, and recent work shows that states 1 to 4 in particular co-activated with memory replay^24^, while most other network states were less likely to be active. Here, we found cycle structure to persist in this dataset (figure 6A; cycle strength, mean (standard deviation): ***S*** = 0.017 (0.017), p < 0.001 versus permutations), and interestingly, that those network states that have previously been shown be positively correlated with memory replay are clustered in the North face of the circle, while the strongest negatively correlated states are on the opposite phase (Figure 6A, polar histogram).

**Figure 6.**
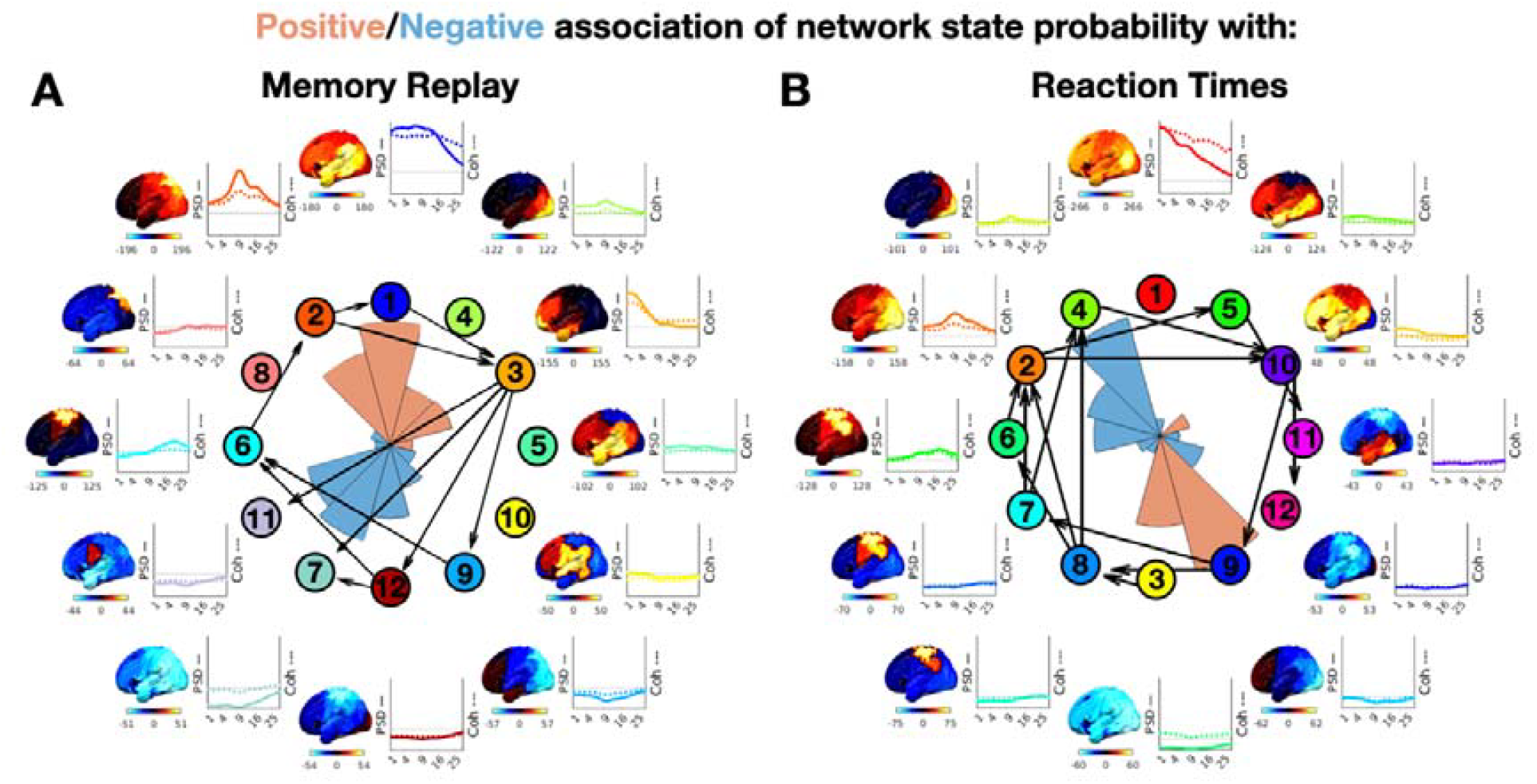
Cycle phase is predictive of cognitive function. Cycle dynamics in a memory replay dataset^24,36^ (A) and a visuo-motor task dataset^37^ (B). Positive/negative associations of network state probability with memory replay (A; ranging from 0-7% increased (red) or decreased (blue) probability) and reaction times (B; correlations ranging from minus (blue) to plus 0.11 (red)) are indicated by polar histogram insets.

These results suggest that these internally generated memory replay events might be precisely timed with respect to the phase of the cyclical activity. However, memory replay does not involve any exogenously prompted behaviour. In particular, it is possible that external events or active behaviour dictate network state activations such that cyclical activity disappears. To answer this question, we applied TINDA to a visual task dataset ^37^.

In the Wakeman-Henson faces dataset ^37^, 19 participants saw a series of famous, unfamiliar, or scrambled faces in six sessions and had to report their asymmetry with a button press. This dataset has previously been shown to elicit task-dependent network dynamics ^38^. Despite the trial structure, we again confirmed the presence of an overall cyclical structure in network state activation (***S*** = 0.058 (0.031), p < 0.001 versus permutations), and we also observed that the ordering of states along the cycle replicated that of the MEG UK dataset (Δθ = 0.84 rad, p = 0.027; Figure 6B). We then correlated the state time courses time locked to button press with the reaction times for each trial. Network probability at 500 ms prior to button onset in each individual was significantly positively correlated with their reaction times in state 3 (R = 0.069, 95% CI = 0.031-0.11, t(18) = 3.8, p=0.0014; t-test against zero), and state 9 (R = 0.11, 95% CI = 0.051-0.17, t(18) = 3.9, p = 0.0009), and negatively correlated with state 2 (R = -0.058, 95% CI = - (0.034-0.083), t(18) = -5.0, p = 0.0001), state 4 (R = - 0.094, 95% CI = - (0.056-0.13), t(18) = -5.2, p = 0.0001), and state 6 (R = -0.051, 95% CI = - (0.023-0.078), t(18) = -3.9, p = 0.001). Notably, as in the replay dataset, the positive and negative correlations respectively clustered on opposite sides of the cycle (Figure 6B, polar histogram). In particular, if there was a high probability that a low power state was active 500 ms before the button press, responses were slower, and vice versa for high power visual/attentional states. Furthermore, when we estimated cycle strength on a trial-by-trial basis (i.e., by running TINDA on the three second segment before a button press), we found a small, but significant Pearson correlation between the cycle strength and reaction times over trials (R = -0.025, 95% CI = - (0.011-0.040), t(18) = -3.8, p = 0.0014), such that higher cycle strength was associated with faster responses. Together, these results indicate that cycle dynamics on a moment-to-moment basis are relevant for cognition.

## Discussion

We show for the first time that the activations of a canonical set of large-scale cortical networks are organised in an inherently cyclical manner, where networks are activated at a preferred phase in a periodic cycle. Furthermore, we show that the cycle’s period, and integrity, relate to age and cognition, while cycle phase is predictive of behaviour on a moment-to-moment basis. Together, these results suggest that cyclical activation of functional brain networks might ensure a periodic activation of essential cognitive functions.

### Organisational structures in functional brain networks

Previous research in fMRI has shown a dissociation of resting state networks (RSNs) into cognitive and perceptual clusters or “meta states” ^14,39–41^. In particular, states within the perceptual/cognitive clusters were highly correlated in terms of temporal occurrence ^14,39^ and connectivity profile ^41^, but not in states between clusters. While fMRI and MEG have different biophysical origins and temporal sensitivity, the spatial extent of RSNs is remarkably similar ^11,17,38^. Our results indeed suggest a dissociation of perceptual and cognitive network states, by positioning them on opposite phases of the cycle, most clearly observed for the default mode network (DMN) and dorsal attention network (DAN), see figure 3. Moreover, it suggests a preferred pathway of state transitions between these extrema.

### Broken detailed balance in brain activity

Network state transition asymmetries like these have further been linked to macroscopic broken detailed balance. This deviation from thermodynamic equilibrium is a hallmark of living systems and can be directly linked to energy consumption and system complexity^42^. Previous research in this field has shown that the level of broken detailed balance correlates with the level of consciousness^20,21^ and cognitive exertion^19,22^, and to show potential as a biomarker for progressive brain disorders^43^. Here, we add novel insights into the fast transient networks of oscillatory power and synchronization underlying macroscale broken detailed balance and reveal the time scale at which these cognitively relevant networks cycle. Furthermore, we found that macroscale broken detailed balance increases with age and is stronger on longer time scales, though it is unknown how different methodological approaches interrelate and provide insights into the temporal sensitivity of broken detailed balance.

### Motifs in brain networks

While prior research has established that transitions between brain network states are not random, identifying phenomena like “asymmetric transitions”, “repeated motifs,” or potentially localized “cyclical motifs” ^44,45^, these findings differ significantly from the “cyclical pattern” investigated in this paper. Previous work typically highlighted specific, often localized, aspects of network dynamics: asymmetric transitions show a preferred direction between two states (A➔B is more likely than B➔A), and repeated or cyclical motifs might reveal recurring short sequences or small loops involving a subset of networks. For example, Sporns and Kötter showed that some motifs are more prevalent in anatomical connectomes than in random networks, some of which are cyclical^44^. However, none of these necessarily imply the existence of a global, overarching cycle that incorporates a full set of canonical networks in a specific, repeatable order. For example, strong asymmetry between a few states or the existence of a small recurring motif, like A➔B➔C➔A (e.g. VP➔V3➔V2➔VP in the macaque visual cortex^46^), does not guarantee that the system tends to progress through all other major network states, e.g. D, E, F, etc., in a consistent sequence before returning. The current study’s “cyclical pattern” posits this more comprehensive, large-scale temporal organization, suggesting that the brain tends to flow through the full set of recognisable, canonical large-scale cortical networks over hundreds of milliseconds, a distinct concept from previously described local transition biases or mini-sequences.

### Time scales of structured brain dynamics

Previous studies investigating the asymmetry in functional brain networks have either focused on Markovian state transitions^11,22^, or the (time-lagged) correlation between network activation patterns ^2,5,14,19^. TINDA differs from these by considering the general pattern in network transitions beyond the direct (i.e., Markovian) transitions. This revealed that asymmetric network transitions occur to a different extent at different time scales, with strongest asymmetries on >2 second time scales (figure 3). These asymmetries described an overall cyclical activation pattern, which, due to the stochasticity of individual cycles, had lower typical durations of 300-1000 ms (figure 5 and Supplement VIII), an order of magnitude larger than the typical lifetime of a cortical network state^11^. These timescales have previously been shown to be a lower limit for scale-free global brain dynamics^1^ and the most relevant timescale for global brain processing^47,48^. While we have shown that cycle dynamics at these temporal scales are relevant for behaviour on a moment-to-moment basis (see Relevance for cognition), future work should further explore their role in different cognitive tasks and at difference temporal scales.

### Cycle dynamics and age

Interestingly, we found these time scales to lengthen with age, concurrent to an increase in cycle strength. This is in line with age-related cognitive decline and slowing^49^, though correlations with cognitive performance indicated a more complex relationship. Other age-dependent changes in brain activity are ubiquitous and include a slowing in the power spectrum^50–52^ and a decrease in network connectivity, which has been related to a decrease in the segregation of functional networks^53–57^.

### Heritability of cycle metrics

Another observation that argues for cyclical dynamics to be rooted in our biology is its strong genetic component of cycle rate. Other heritable components to large-scale brain networks have been shown in the past, including connectivity in specific functional networks ^14,58–61^, and frequency bands ^62^, and static connectivity ^63^. In particular, Vidaurre et al. found that the degree to which an individual spent more time in either perceptual or cognitive fMRI resting states was heritable^14^ . How these and other fMRI dynamics are related to the cyclical dynamics described here is a topic for future research.

### Relevance for cognition

While most datasets that we explored here involved wakeful rest, the cyclical dynamics also persisted in task data. Moreover, the phase within the cycle and cycle strength were predictive of cognitive function. While the HMM framework has successfully shown large-scale cortical network associations with cognitive function before^24,38^, here we add that positive and negative associations with memory replay, or response speed, were predicted by network states on oppositive phases of the cycle. One question that arises is whether cycle dynamics like speed and phase can be (consciously) controlled or disrupted by a cognitive task, which is expected from the stochasticity of state transitions within the cycle. On the other hand, the persistence of the ordering of network states within the cycle and the detrimental effect of cycle phase on certain cognitive functions suggests it could reflect a homeostatic process. In fact, homeostatic cyclical rhythms are omnipresent in biological systems^64^, with the sleep cycle as one of the most well-known examples^65^. In sleep, cycling through each of the five functional stages allows the body to experience the benefits of each stage multiple times throughout the night, ensuring each function is carried out even if sleep is disrupted. Similarly, cycles in large-scale brain networks could ensure periodic activation of essential cognitive functions, with stochasticity enabling cognitive flexibility.

### Limitations and Future Directions

The current study comes with a number of limitations. Firstly, the TINDA method is a post-hoc analysis tool that is used on binarised state time courses (i.e., brain networks are either “on” or “off”), and furthermore, it does not incorporate an explicit model of long-term (variable time) state transitions. In future work, we hope to deploy non-Markovian models like neural networks for inferring brain networks (such as DyNeMo^66^), but it remains an open question how to adapt TINDA to state time courses that are not mutually exclusive. While we have reproduced our main results in multiple datasets, some results could not be reproduced, i.e., the heritability of cycle metrics and the association of cycle metrics with cognitive scores. Replicating these analyses in independent datasets is essential but rely on the availability of the relevant data. This would also clarify the role of cycle dynamics for cognition across individuals and their potential as biomarkers for disease.

Another limitation of the current study and the field of functional brain networks more generally, is a lack of taxonomy with respect to the macro scale functional brain networks. This can lead to ambiguity or overinterpretation of the functional network, and it is unclear in what capacity they are rooted in the underlying physiology ^67^. Moreover, there is no consensus in electrophysiology about which features constitute a brain network, be it coherence, power, spectral shape, et cetera, and how to relate these to brain networks observed in fMRI. Regarding the first point, we argue that a principled definition of a brain network is one where networks can be distinguished, not by a single arbitrarily chosen feature, but instead by multiple network features. We therefore use the time-delay embedded (TDE) HMM ^29^, in which brain networks are characterised by distinct auto- and cross spectral properties as part of a generative model that is capable of explaining the full signal content. Previous work has also shown that TDE-HMM results in identifiable networks that are high reproducible across different data, sites, and task/rest designs, which validates this approach. Secondly, there is growing effort of comparing functional brain networks across studies ^29,68,69^, and modalities ^70–76^. We have made quantitative comparisons of MEG state topographies with the widely used fMRI based Yeo7 parcellation ^31^ (see Supplement XII), from which we tentatively concluded that the cycle separates the default mode network (top of the cycle) and the dorsal attention network (bottom of the cycle). However, we do note that the existence and presence of cycles shown here in five independent datasets do not rely on the physiological interpretation of individual networks. More efforts need to be made in quantitatively comparing functional brain networks inferred from electrophysiology and hemodynamic responses, particularly in simultaneous recordings.

## Methods

All analyses were carried out in MATLAB ^77^ and Python, using in-house developed software packages OHBA Software Library ^78–80^, HMM-MAR^81^, OSL-dynamics ^69^, and MNE-Python^82,83^. The TINDA package is available for MATLAB ^84^, and Python ^85^.

### Data

We used data from five MEG datasets: Nottingham MEG UK (N = 55), Cam-CAN (N = 612), HCP (N = 79), Replay (N=43), and Wakeman-Henson (N=19). MEG UK, Cam-CAN, HCP, and replay contain MEG resting state data, Replay and Wakeman-Henson MEG task data, and all but the replay dataset include T1 weighted MRI brain scans. Datasets include demographic data and in the case of HCP, heritability data. Ethics and consent details are outlined separately for each of these datasets below.

#### MEG UK

The UK MEG Partnership data comprised 77 healthy participants recruited at the University of Nottingham, of which 55 were used after discarding 22 subjects because of excessive head movements or artefacts. The dataset contains structural MRI scans and MEG data from a CTF MEG system containing 275 axial gradiometers with a sampling frequency of 1200 Hz. The participant group had a mean age of 26.5 years (range 18-48). 20 of them were female, and 35 of them were male. All participants gave written informed consent, and ethical approval was granted by the University of Nottingham Medical School Research Ethics Committee. The MEG data comprised roughly 5 to 6 minutes eyes-open resting state and has previously been used to characterise MEG resting state network dynamics ^24,29,86^.

#### Cam-CAN

The Cambridge Centre for Aging Neuroscience (Cam-CAN) dataset comprised data of 700 healthy participants recruited at the University of Cambridge, of which 612 were used here. The dataset contains structural MRI scans and MEG data from an Elekta Neuromag Vectorview system containing 102 magnetometers and 204 orthogonal planar gradiometers, with a sampling rate of 1 kHz. The participants were aged 18-88, with 83-95 participants per age decile (except in the 18-28 decile, which counts 45), 310 were male and 302 were female, equally distributed across the age deciles. All participants gave written informed consent, and ethical approval was granted by the Cambridgeshire Research Ethics Committee. The MEG data comprised of approximately 9 minutes eyes-closed resting state.

#### HCP

The MEG component of the Human Connectome Project (HCP) comprised 100 healthy participants recruited at the Saint Louis University, of which 79 were used after discarding subjects with excessive variance. The dataset contains structural MRI scans and MEG data from a 4D Neuroimaging MAGNES 3600 MEG system containing 248 magnetometers sampled at 2 kHz. The participant group had a mean age of 29 (range 22-35) of which 37 females and 42 males and contained data of 13 monozygotic twin pairs and 11 dizygotic twin pairs. All participants gave written informed consent, and ethical approval was granted by the local ethics committee. The MEG data comprised of three times 6 minutes of eyes-open resting state.

#### Replay

The Replay data^36^ contained a primary dataset (dataset 1), and a replication dataset (dataset 2). For both datasets, participants were scanned on a 275 channel CTF MEG system while engaged in a localiser task, a sequence learning task, and periods of rest. Activations corresponding to images in the localiser task were found to replay during rest, in the sequence that corresponded to the learned sequences. The top 1% replay probabilities were here defined as the memory replay events, as in Higgins et al^24^. Replay dataset 1, was acquired from 25 participants with a mean age of 24.9 (range 19-34) of which 11 males and 14 females. Four subjects were excluded due to large motion artifacts or missing trigger information. All participants signed written consent in advance; ethical approval for the experiment was obtained from the Research Ethics Committee at University College London under ethics number 9929/002. Replay dataset 2 was acquired from 26 participants with a mean age of 25.5 (range 19-34) of which 10 males and 16 females. Four participants were later excluded due to motion artifacts or failure to complete the task. All participants signed written consent in advance; ethical approval for the experiment was obtained from the Research Ethics Committee at University College London under ethics number 9929/002. In the current study, replay datasets 1 and 2 were analysed jointly.

#### Wakeman-Henson

The Wakeman-Henson faces dataset^37^ comprised MEG data acquired on an Elekta Neuromag Vectorview system of 19 participants. Of these, 8 were female, and 11 were male, and the sage range was 23-37 years. All participants gave written informed consent, and ethical approval was obtained from the Cambridge University Psychological Ethics Committee. Each participant completed six sessions of a perceptual task in which they would see a famous, familiar, or scrambled face, to which they had to respond based on the symmetry of the image. Each trial begins with a fixation cross onset between 400 and 600 ms before a target stimulus appears. The target is either the face or scrambled face stimulus and remains onscreen for between 800 and 1000 ms. Further details can be found in ^37^.

### Preprocessing

MEG data were co-registered to the MRI structural scans, or to fiducial markers in the Replay data where MRI structural scans were not available. The MEG UK and Cam-CAN data were downsampled to 250 Hz, filtered in the 1-45 Hz range (using zero-phase digital filtering so that effects are symmetrical across time), and source-reconstructed using an LCMV beamformer to 3559 dipoles. The dipoles were then combined into 38 parcels spanning the entire cortex by taking the first principal component of all dipoles in a parcel. This parcellation was used previously to estimate large-scale static functional connectivity networks in MEG ^50^. The HCP data were downsampled to 240 Hz, filtered in the 1-80 Hz range, and source-reconstructed using an LCMV beamformer to 5798 dipoles. The dipoles were then combined into 78 parcels of the AAL parcellation^87^ spanning the entire cortex by taking the first principal component of all dipoles in a parcel. Bad segments were removed manually and correction for spatial leakage was applied using symmetric multivariate leakage correction ^88^. Finally, the potential inconsistency over subjects of ambiguous source polarity was removed using sign-flipping based on lagged partial correlations ^38^.

### Hidden Markov modelling

To find large scale brain networks in a data driven way, we applied a time delay embedded hidden Markov model (TDE-HMM) with 12 states and 15 embeddings, corresponding to lags of -28 to +28 ms (-29 to +29 ms for HCP). Note that we refer to the HMM states as “network states” to reflect that the method is designed, and has been shown, to find states that represent distinct cortical networks of oscillatory brain activity in MEG/EEG data^29^ . The HMM framework is a generative model that assumes there exist a finite number (***K***) of recurring, transient, and mutually exclusive hidden states that generate the observed data. Here, each state is characterised by a spatio-spectral profile (i.e., in terms of PSD and connectivity in/across regions). Thus, every time point in the data is associated with one of the states [**1**: **2**, … ***K***], and the sequence of states is assumed to be Markovian. This means that the state active at time point ***t*** only depends on the state active at ***t* − 1**, captured by the transition probability between all states. We used a multivariate Gaussian observation model with zero mean. Models were inferred separately for the MEG UK, Cam-CAN, and HCP datasets. For the Replay datasets, we kept the model from the MEG UK dataset fixed, and subsequently fitted it to the replay data, as in Higgins et al^24^.

### Spectral analysis

We estimated the spectral information (PSD and coherence) for each state by fitting a multitaper to the original, parcellated data, condition on the active functional brain network, as in Vidaurre et al.^29^ The multitaper used a taper window length of 2 s, a frequency range of 1 to 45 Hz with a 0.5 Hz resolution (i.e., applying 7 Slepian tapers). This reflects the full multivariate model parameter space (an array that is frequencies x channels x channels x states), however the high model dimensionality necessitates further dimensionality reduction methods if this information is to be visualised. We reduce the spectral dimensionality using spectral mode decomposition, resulting in spatial power and coherence maps for a data-driven set of frequency band modes. This decomposition is implemented by non-negative matrix factorization^29,38^. We fitted this with two modes to separate wideband activity from high frequency noise (Figure S1). We then used the wideband mode to weight the frequencies of the individual states when producing topographies.

### Ordering the HMM states

We ordered the HMM states based on state coherence using the MEG UK dataset. States inferred from the MEGUK dataset were reordered based on the mean coherence in that state, from high (state 1) to low (state 12) coherence. The orderings in the other datasets (Cam-CAN/HCP/Wakeman-Henson) were then matched to the MEG UK ordering as follows. First, the correlation was computed between the coherence of each pair of states (where a “pair of states” consisted of one state from MEGUK and one state from (e.g.) Cam-CAN. The correlations were then used as a cost function to solve the linear assignment problem using the *matchpairs* function in MATLAB^89^, matching every state in the (e.g.) Cam-CAN dataset to a state in the MEG UK dataset. Due to the different parcellation used in the HCP dataset, here we used the correlation between power maps in MNI volume space as cost function. In figures throughout the manuscript, state numbers thus indicate equivalent (i.e., “best matching”) states, whereas state colours were different between datasets, to stress that state descriptions were inferred independently for each dataset.

### Temporal Interval Network Density Analysis (TINDA)

We developed the TINDA method to analyse inter-state dynamics in the context of dispersive inter-state intervals (ISI’s). We first partition all observed interstate intervals, defining *T*_1_^*m,i*^ to be the set of timepoints that fall within the first half of ISI’s for state m; and *T*_2_^*m,i*^ to be the set of all timepoints that fall within the second half of these ISI’s. We then compute the ***K*** × ***K*** fractional occupancy asymmetry matrix (***A***), defined as the matrix whose (m,n)-th entry is:

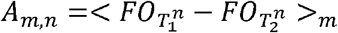

Where <…> _*m*_ denotes the average FO difference for state *n* over state *m* ISI’s (Figure 1). These Fractional Occupancy Asymmetry matrices are computed for each subject.

### Cycle detection and Cycle Strength

TINDA establishes whether there is a general flow of states into and out of a particular state. We investigated whether this pattern is embedded in a larger, hierarchical structure, specifically a cycle. We interpret the fractional occupancy asymmetry matrix, A, as a weighted, directed graph of ***K*** nodes (i.e., number of states) and ***K***^**2**^ − ***K*** edges (i.e., from every node to every other node). The fractional occupancy asymmetry thus defines the weight and direction of each edge.

To investigate how these edges relate to specifically cyclical dynamics, we define a metric of cycle strength for each configuration of the ***K*** nodes around the unit circle. Each node is associated with a phase *q*, positioned on the unit circle in 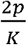 intervals, spanning [0, 2*p*]. We can then represent each directed transition, from state n to state m, by a vector in the complex plane defined by the phase difference between the relative position of nodes m and n:

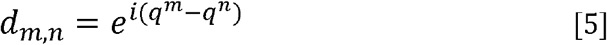

The magnitude of the imaginary component of this vector represents a geometric projection of each state transition onto the plane tangential to the unit circle at node n. Trivially, the cumulative sum of these vectors for all ***n*** and ***m*** is zero. However, if these vectors are weighted by the strength (and direction) of the corresponding FO asymmetry, then the sum of their imaginary components represents the cumulative strength (i.e., of the asymmetry) and the polarity represents the net direction (i.e., clockwise (+) vs. counterclockwise (-)) of transitions tangential to the unit circle. Hence, we define the cycle strength, S, is defined as:

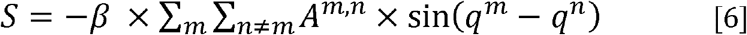

Where *β* is a normalisation factor based on the theoretical maximum cycle strength for ***K*** states, such that ***S*** is constrained to be [-1, 1]. The theoretical maximum cycle strength is computed for K states by assuming a perfect asymmetry of +1 for all possible clockwise connections, and -1 for all possible negative connections.

We permuted the position of each state on the unit circle (i.e., the node identity) to maximise ***S***. This reveals the sequence of states for which the overall directionality is maximal in the clockwise direction. Note that we could have chosen to maximise negative ***S*** instead. This would have only resulted in all circle plots going in a counterclockwise direction; it would not have changed any of our results.

### Circle plots

The circle plots all show the network states in the sequence which maximises the cycle strength (in clockwise direction). The edges ***E*** that are shown are those where the fractional occupancy asymmetry is statistically significant (see Statistics), where the direction of the arrow depends on the sign of the corresponding edge asymmetry, where ***α*** is the (corrected) statistical threshold (see Statistics):

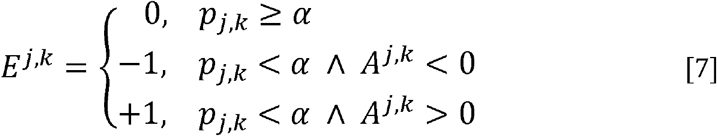

### Cycle rate

To quantify cycle rate, we applied a post-hoc analysis to the state time course parameters already learned. Specifically, we derived a feature from the state time courses defined as the number of state visits in a sliding window equal to the average state lifetime (64-68 ms, depending on the dataset). We then fit a second level HMM to this feature time course, where this HMM used a Poisson observation model and sequential Markov dynamics ^90^. We selected a model with ***K*** = **4** states, where we initialised the state probabilities as the distance (i.e., in circle space) to the centroid of each of the four modes in figure 4B-C. We also enforced a sequence of state 1 > state 2 > state 3 > state 4 > state 1, etc. such that a single cycle was defined as sequentially activation each of the four modes in Figure 4B-C. Using this initialised model, we inferred the state time courses from the data without training the model to convergence. This was done to not deviate from our definition of a “cycle”, and to subsequently quantify cycle duration as the time it takes to cycle through a full 1 > 2 > 3 > 4 sequence of the second level HMM. For correlations with individual traits, the inverse (i.e., cycle rate) was used, as this was more normally distributed.

### Statistics

#### FO asymmetry and Circle Plots

Circle plots show the edges where the FO asymmetry is strongest (and significant). In order to test for significance, the FO Asymmetry was tested on the group level with a two-tailed dependent samples T-test for each of the connections ***m, n***, where the alpha threshold of 0.05 was Bonferroni corrected for 132 tests (i.e., ***K***^**2**^ − ***K***), resulting in a corrected alpha threshold of 0.00038. Due to the large subject numbers in HCP and Cam-CAN, more stringent thresholds were applied in these datasets. For Cam-CAN, the edges with absolute t-values larger than 11 are shown (corresponding to a p-value smaller than 3.8*10^-24^), and for the HCP dataset the edges with absolute t-values larger than 4.3 (corresponding to p-values smaller than 3.8*10^-5^).

#### Cycle strength

We report the results for the sequence of network states that results in the strongest, clockwise cycles. A claim of nonzero cycle strength could be a trivial consequence of this optimisation. For this reason, we compared the observed cycle strength with that from permutations, where for each permutation we permuted each subjects’ state labels and recomputed the FO asymmetry and optimised state ordering. This was done 1000 times. The observed cycle strength was compared with the permuted versions at alpha=0.05.

#### Within-subject consistency of cycle metrics

The individual consistency of cycle metrics was directly estimated using the intraclass correlation efficient (ICC) in MATLAB, with type ‘1-1’ as implemented in Salarian (2024)^91^.

#### Correlation with individual traits

For the correlation of cycle rate and cycle strength with individual traits, we first regressed out heart rate. Outliers more than three standard deviations from the mean value were removed, and cycle metrics were normalised, prior to the general linear model (GLM). We fitted a mean term, and the cycle rate and cycle strength to age and sex separately, using a Gaussian and binomial distribution, respectively. Beta terms for each were significant if the corresponding p-value was lower than the alpha threshold of 0.0125 (i.e., 0.05 corrected for 4 tests).

#### Heritability

In order to test whether variance in cycle metric could be explained by genetic factors, we used an ACE model, as implemented in the Accelerated Permutation Inference for the ACE Model (APACE) framework ^92^. APACE was run on all subjects’ cycle metrics for the three resting state sessions, separately for cycle rate, and cycle strength, using 10000 permutations. Alpha thresholds of 0.05 were Bonferroni corrected for two tests. To ensure estimated heritability effect were not caused by common demographic and morphometric measures, we repeated the analysis after regressing out the following confounds in stepwise fashion (See Supplement IX): age, the square of age, sex, an age and sex interaction, an interaction between sex and the square of age, the cube root of intra-cranial volume and of cortical volume (both estimated with FreeSurfer^93^).

#### Correlation with cognitive scores

A canonical correlation analysis (CCA) was executed on the Cam-CAN dataset, between the cyclical summary metrics (cycle rate and strength) on the one hand, and thirteen cognitive scores on the other hand. For all metrics, we first regressed out age, and sex, and heart rate, and then z-transformed the data. The CCA resulted in two CCA components, which were tested for significance by comparing against a permutation distribution of 10000 permutations, where for each permutation, the cognitive scores were shuffled over subjects.

#### Correlations with reaction times in Wakeman-Henson

We time locked the state probability time courses to the button presses in the Wakeman-Henson data and correlated state probability at 500 ms prior to the button press with the reaction time on that trial. This was done separately for each session and state, after which we averaged the correlations over sessions for each subject. We tested whether the correlation was significantly different from zero for each state with a paired t-test, using a Bonferroni corrected alpha level of 0.05/K. Similarly, we correlated reaction times with an instantaneous estimate of cycle strength, computed by running TINDA on 3 second segments prior to the button press, and calculating the cycle strength. Correlations were tested against zero on the group-level using a paired t-test.

## Data and Code Availability

The code for all analysis code described here is publicly available and can be accessed at https://github.com/OHBA-analysis/Tinda. The MEG UK Partnership data is held by the MEG UK Partnership, for which access can be requested at https://meguk.ac.uk/database/. The Cam-CAN dataset is available upon request to https://camcan-archive.mrc-cbu.cam.ac.uk/dataaccess/datarequest.php. The HCP dataset is freely available at https://db.humanconnectome.org/app/template/Login.vm but will require an application for sensitive data (see https://www.humanconnectome.org/storage/app/media/documentation/LS2.0/LS_Release_2.0_Access_Instructions_June2022.pdf). The Replay dataset will be freely available upon request (subject to participant consent) to. The Wakeman-Henson dataset is publicly available at OpenNeuro (https://openneuro.org/datasets/ds000117/versions/1.0.5).

## Acknowledgements

The authors would like to thank Thomas Nichols and Xu Chen for their help in using the APACE model. MWW’s research is supported by the Wellcome Trust (106183/Z/14/Z, 215573/Z/19/Z), the New Therapeutics in Alzheimer’s Diseases (NTAD) and Synaptic Health in Neurodegeneration (SHINE) studies supported by UK MRC and the Dementia Platform UK (RG94383/RG89702), and the NIHR Oxford Health Biomedical Research Centre (NIHR203316). DV is supported by a Novo Nordisk Emerging Investigator Award (NNF19OC-0054895) and by the European Research Council (ERC-StG-2019-850404). The Wellcome Centre for Integrative Neuroimaging is supported by core funding from the Wellcome Trust (203139/Z/16/Z and 203139/A/16/Z). For the purpose of open access, the author has applied a CC BY public copyright licence to any Author Accepted Manuscript version arising from this submission. The views expressed are those of the author(s) and not necessarily those of the NIHR or the Department of Health and Social Care.

## Author contributions

MVE and CH have contributed equally to the current work and have the right to list their name first when referencing the work. Following CRediT, the following authors contributed to each of the roles: Conceptualization (MVE, CH, MW), Methodology (MVE, CH, CG, AQ, DV, MW), Software (MVE, CH), Validation (MVE, CH), Formal Analysis (MVE, CH, CG, AQ), Investigation (MVE, CH, CG, AQ, DV, MW), Resources (MVE, CH, CG, AQ, DV, MW), Data curation (MVE, CH), Writing – original draft (MVE, CH), Writing – review & editing (MVE, CH, CG, AQ, DV, MW), Visualization (MVE, CH), Supervision (MW), Project administration (MVE, CH), Funding acquisition (DV, MW). All Authors contributed to the article and approved the submitted version.

