## Supplementary Information for "Large-scale cortical networks are organised in structured cycles"

### Supplemental information

#### Supplement 1. Spectral characteristics

We used a non-negative matrix factorization on the complex spectra to separate wide band activity from high frequency noise. Here, we show the wideband components for all studies, as well as the individual state descriptions. Note that the states for the Cam-CAN, HCP, and Wakeman-Henson datasets were reordered to as closely correspond to the state descriptions of the MEG UK dataset (see Methods).

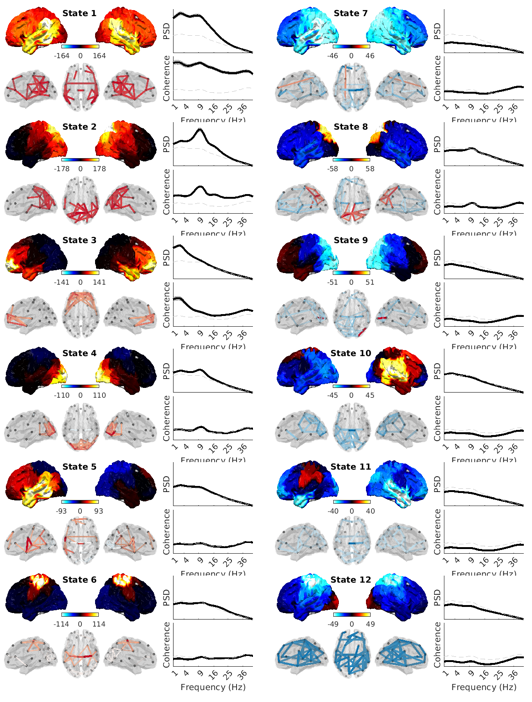

MEG UK

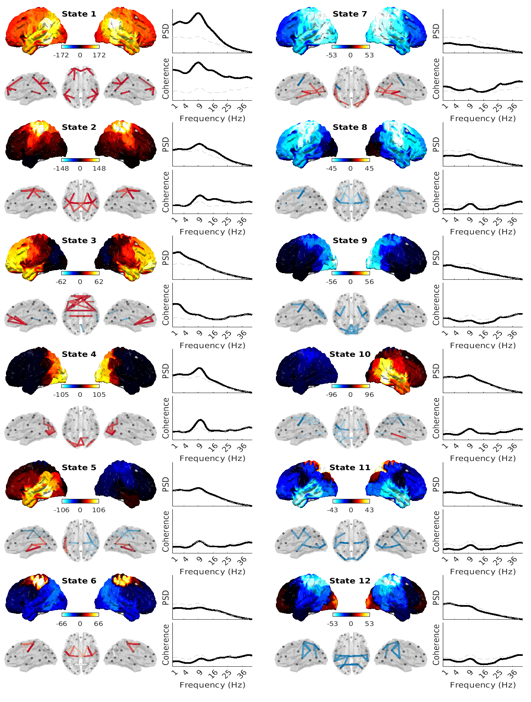

Cam-CAN

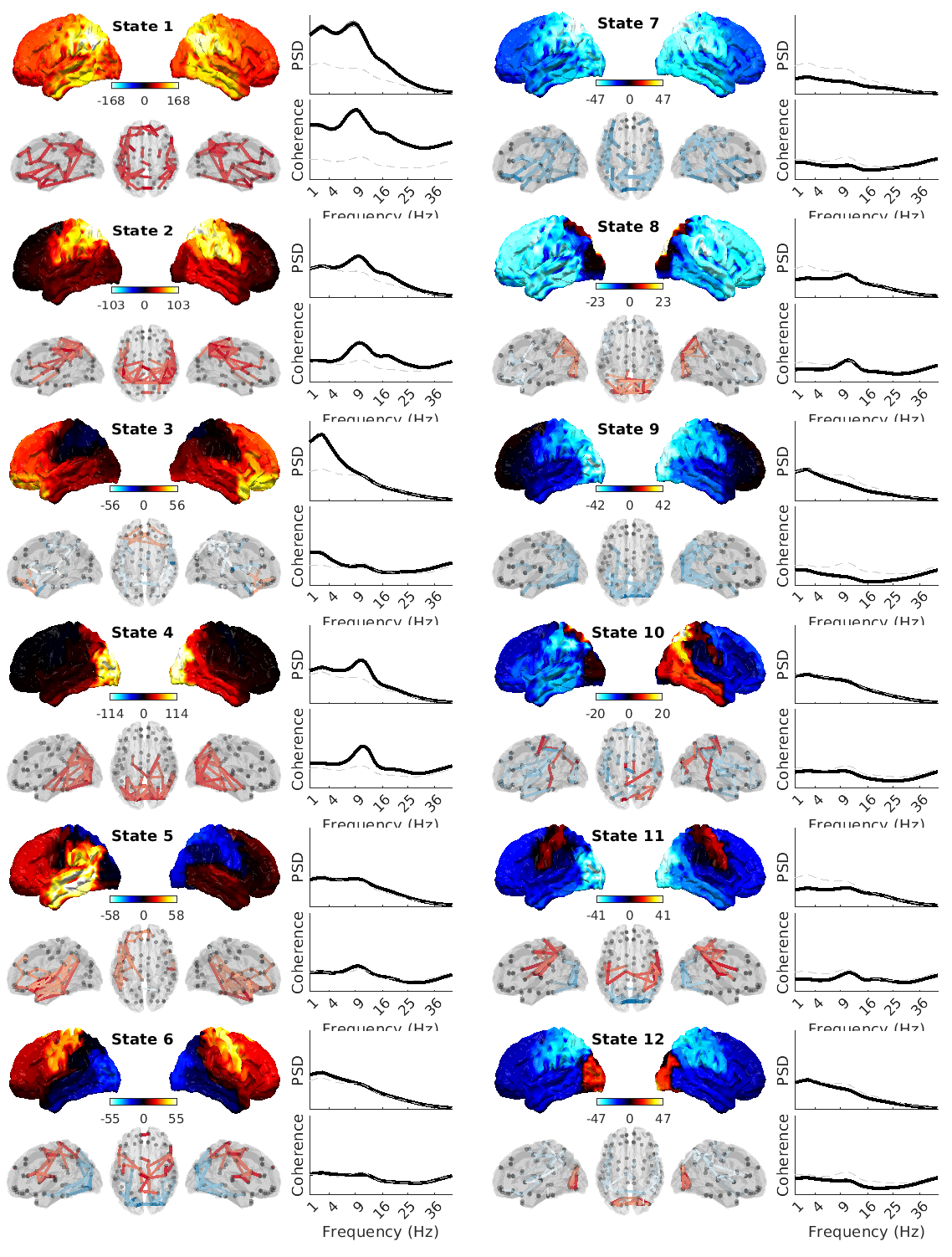

HCP

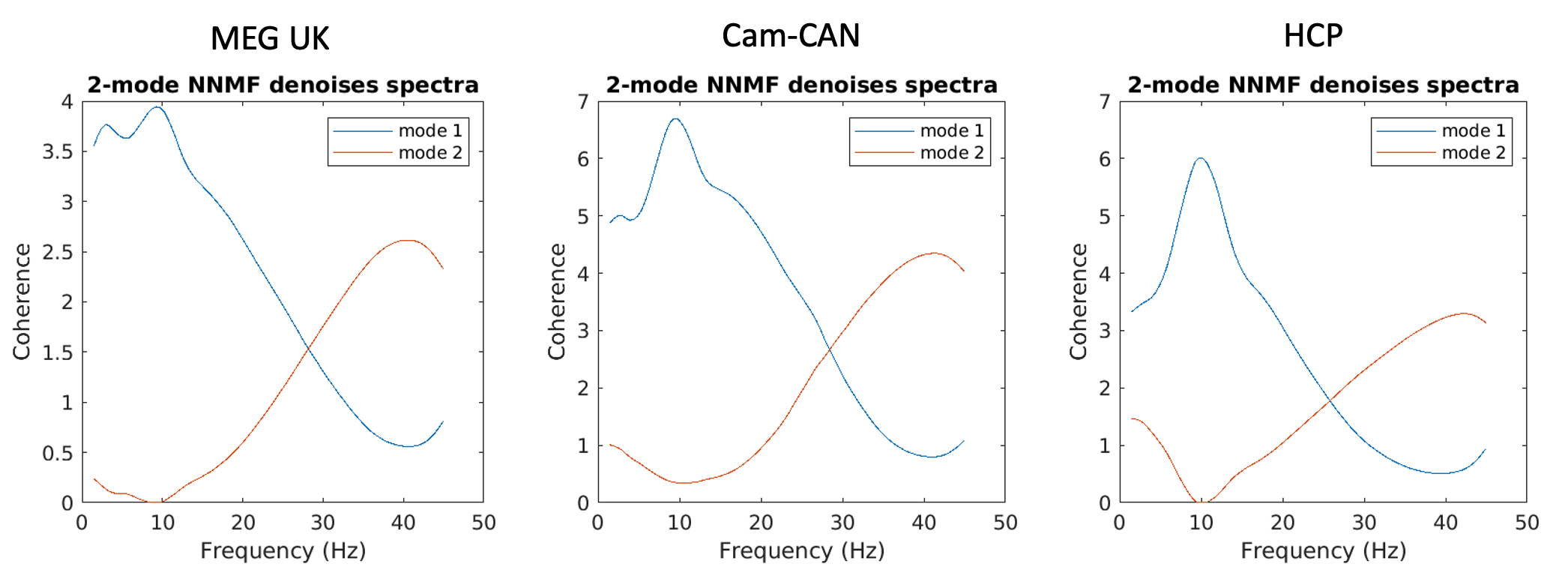

MEG UK

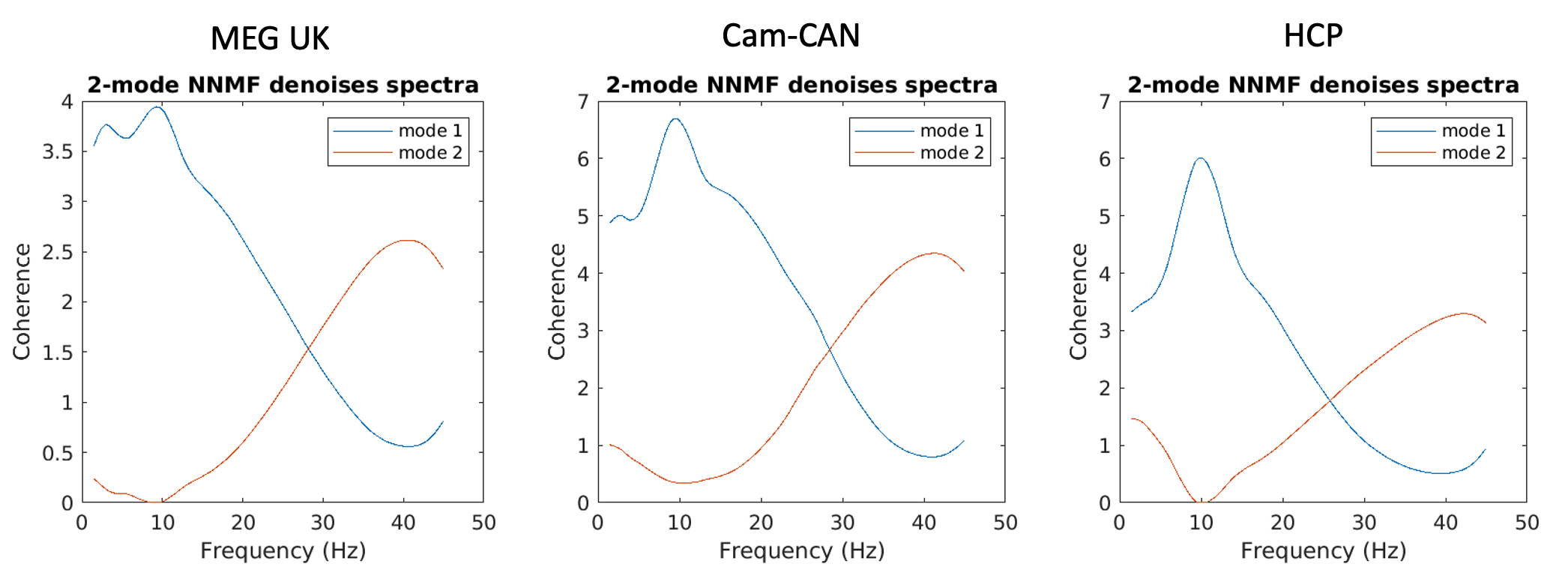

Cam-CAN

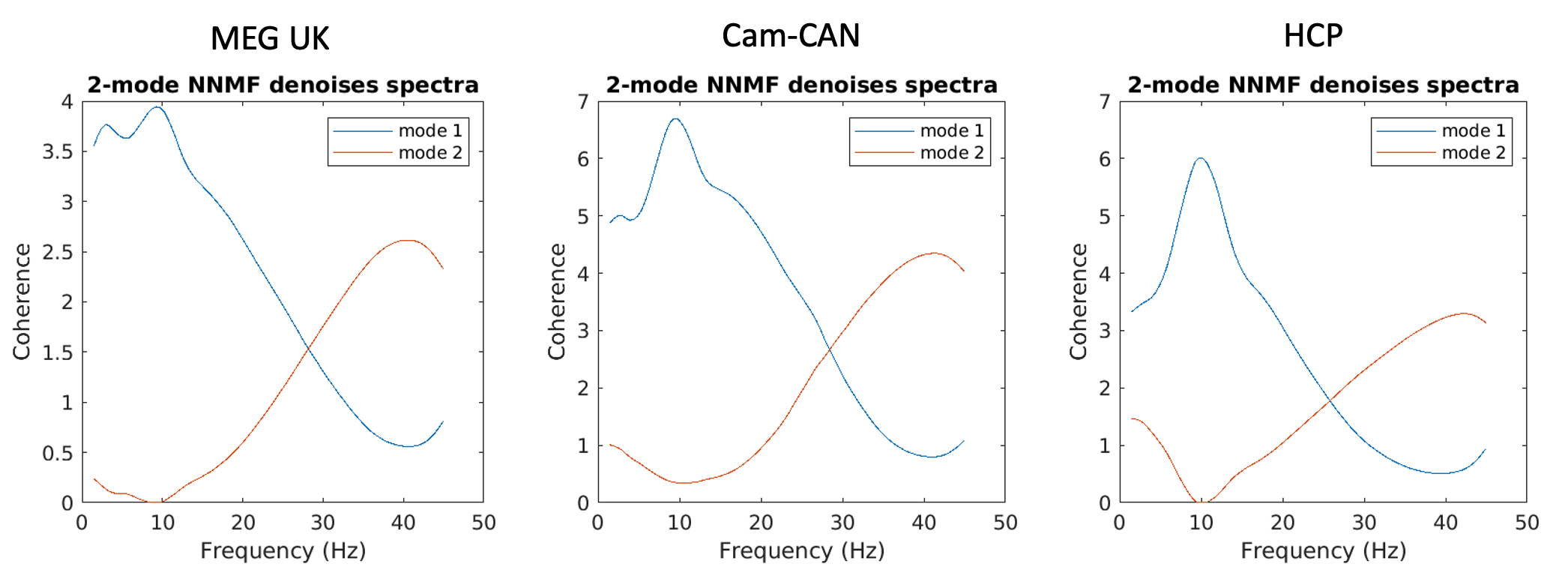

HCP

Wakeman-Henson

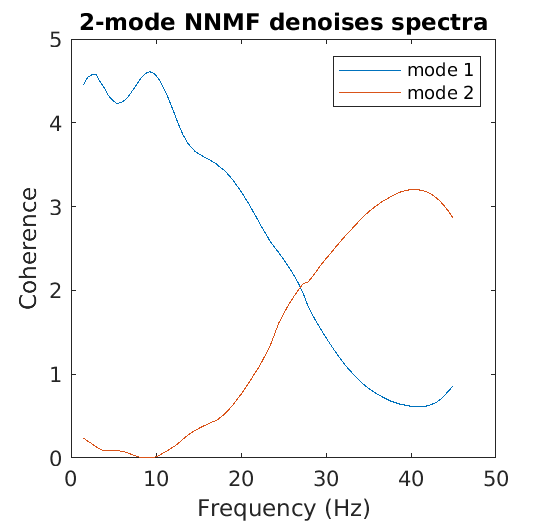

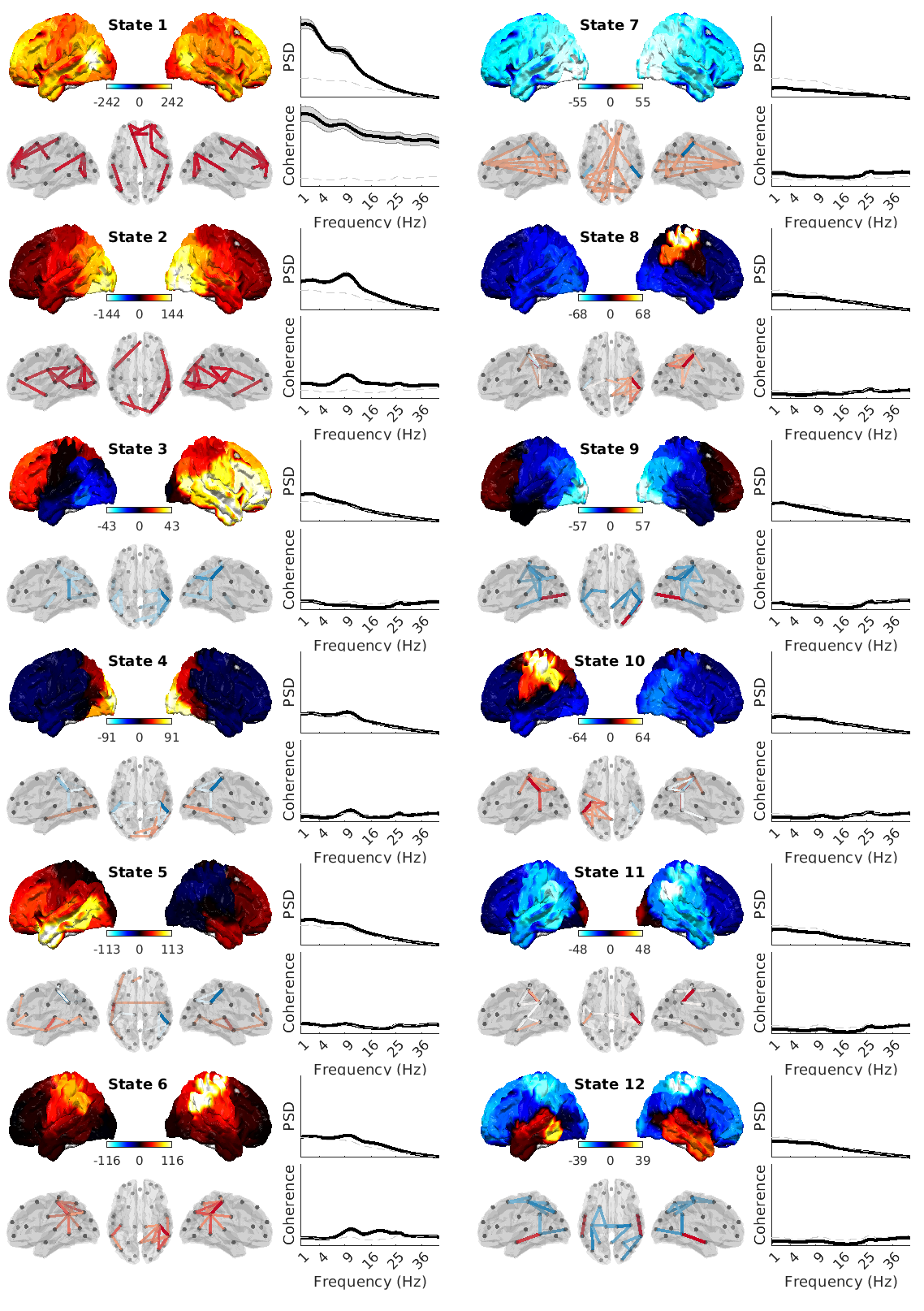

Wakeman-Henson

A

B

Figure S1. Spatial and spectral descriptions of the HMM states. A) The spectral content of each HMM state was summarised in a wideband mode (mode 1; blue) and a high frequency noise mode (mode 2; red) using spectral mode decomposition, in each of the four datasets. Only the wide-band modes were analysed in all further analyses. B) The power/coherence spectra and topographies for all HMM states of the MEG UK, Cam-CAN, HCP, and Wakeman-Henson datasets. For each state, we show the topography of the wideband spectral mode relative to the mean over states (top left; bright blue colours indicate strong reductions in power, bright red strong increases in power), the power spectrum (top right) of the state (solid line) and mean over states (dotted line), the edges for which the wideband coherence is in the top 2% (5% for MEGUK) relative to the mean over states (bottom left; blue indicate reductions, red increases in coherence), and the coherence spectrum (bottom right) of the state (solid line) and the mean over states (dotted line).

#### Supplement II. Resting state networks have unique and specific transition pathways that do not trivially form cyclical dynamics

In figure 1, we show in the MEG UK data that there are particular states that are more likely to activate before, or after state 1. Here, we show the same for the equivalent state in Cam-CAN (B), and HCP (C). Moreover, we find unique ascending and descending pathways for each state in all datasets (D). Nonetheless, the mere existence of these pathways is not sufficient for cyclical dynamics to emerge, which additionally requires a specific second-order configuration connecting successive pathways (E, p<0.01). To show this, we permuted 100 times the non-diagonal entries of the Fractional Occupancy (FO) Asymmetry matrix, such that the number and strength of ascending and descending pathways were preserved at each node, whilst the global architecture connecting successive pathways was randomised. For each permutation the ordering of states was then optimised to maximise the cycle strength that could arise by chance; no single permutation reached a cycle strength comparable to that observed in real data (p<0.01).

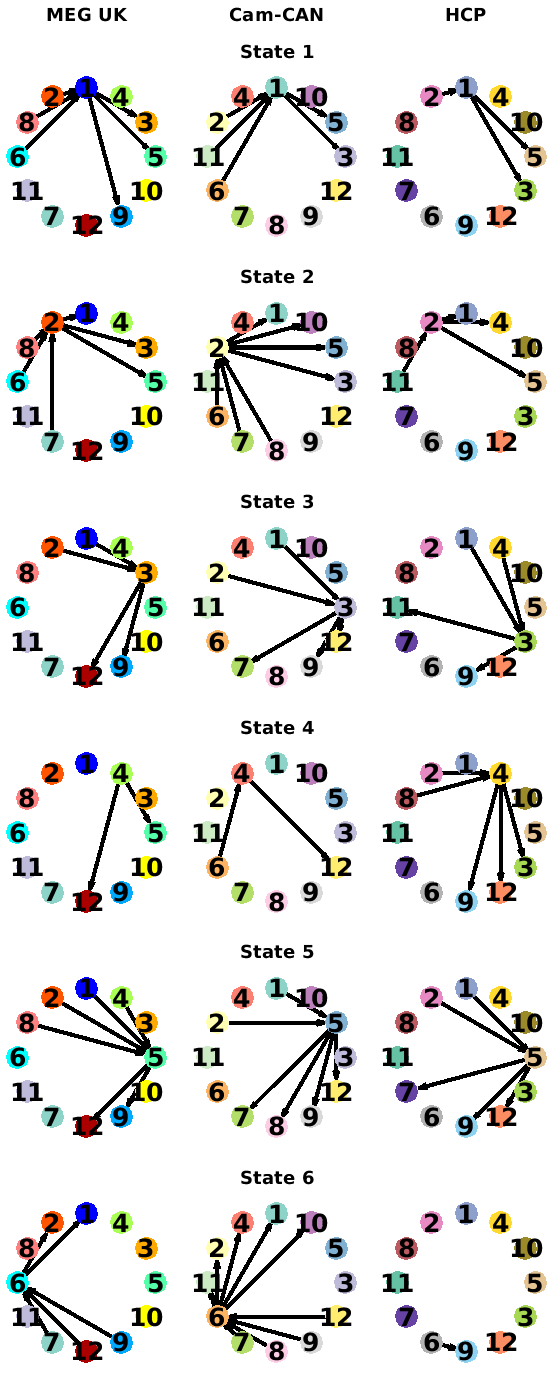

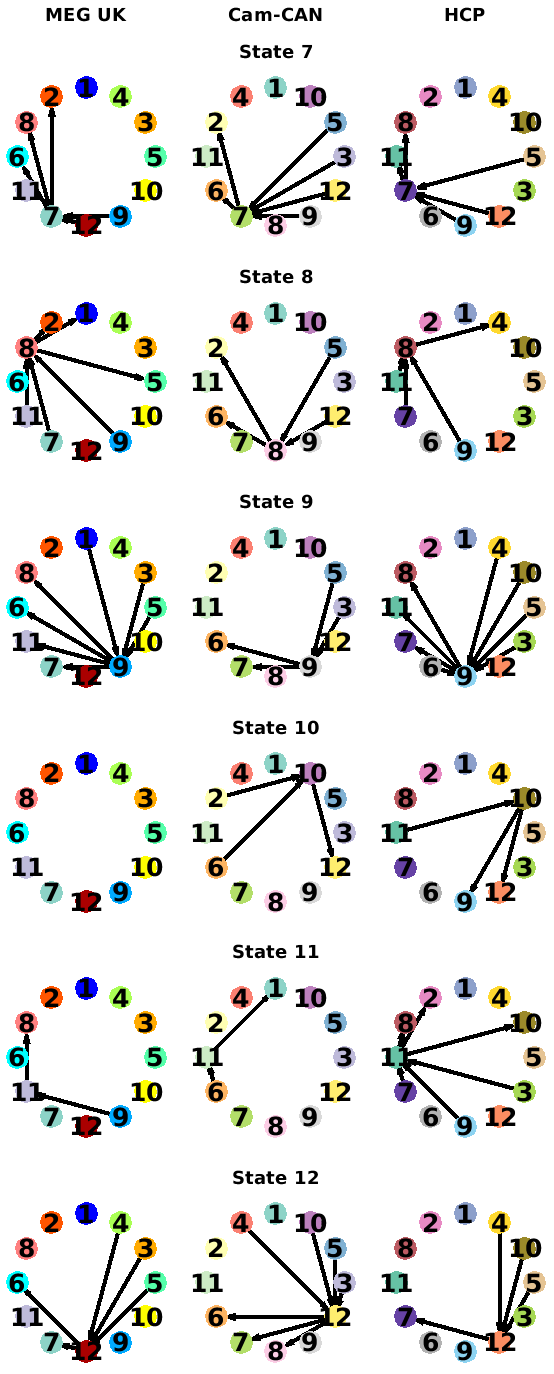

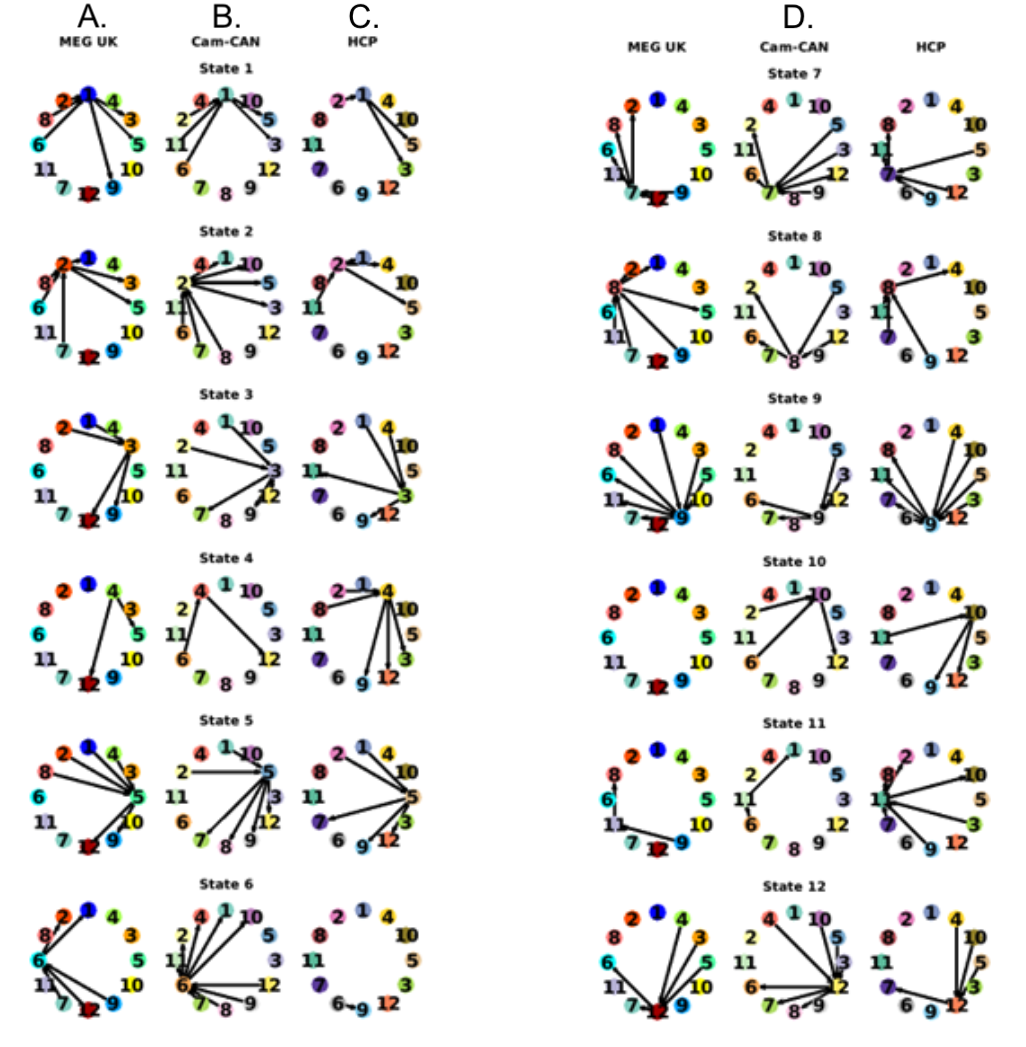

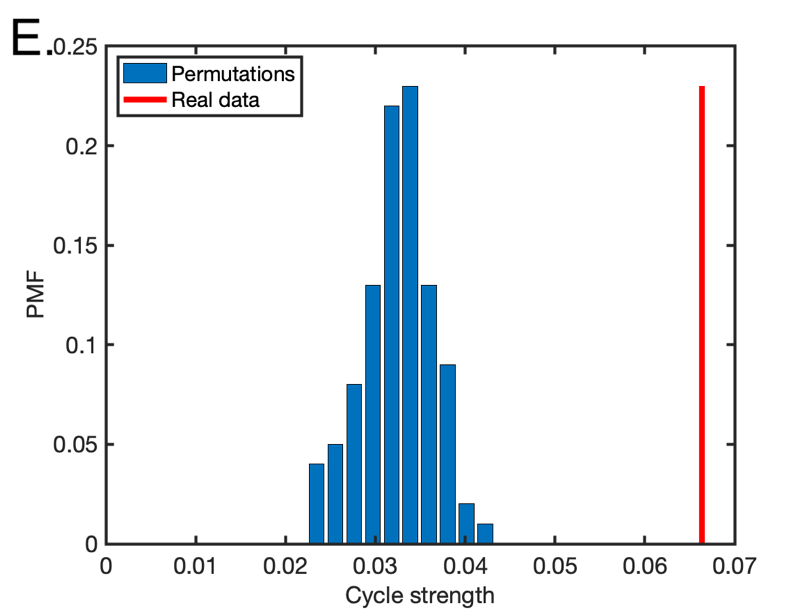

Figure S2. A-C) There are unique transition pathways into, and away from, state 1 in the Nottingham MEGUK dataset (N =55; A), Cam-CAN dataset (N=612; B) and HCP dataset (N=79; C). All arrows shown correspond to significant FO asymmetry between the first and second half of state 1 intervals. Black arrows have a higher FO in the first half of the intervals (i.e., those states activate shortly after the state 1), grey arrows in the second half of the intervals (i.e., those states activate shortly before the next activation of state 1). D) Same as (A-C) but for all other states. E) Cycle strength for the observed Fractional Occupancy Asymmetry, and for the permutation distribution, where in each permutation the off-diagonal entries of the matrix were permuted.

Supplement III. The ordering of states in the cycle is equivalent in the three datasets.

In the main manuscript, we showed that the ordering of states across the cycle is reproduced across datasets. In figure 2, we visualised the optimal ordering of states independently in each dataset (shown here in panels A/B/E). Here, we show how these plots would look if visualising them according to the ordering that was optimised on the first dataset (MEG UK, panels C/F). The different options for ordering the states look qualitatively very similar for both the Cam-CAN and HCP dataset. Quantitatively too, the deviation in cycle strength is minimal when switching between the ordering optimised for the MEG UK dataset or the dataset in question (D/H). These results further substantiate the claim that the order of states in the cycle reproduces across independent datasets.

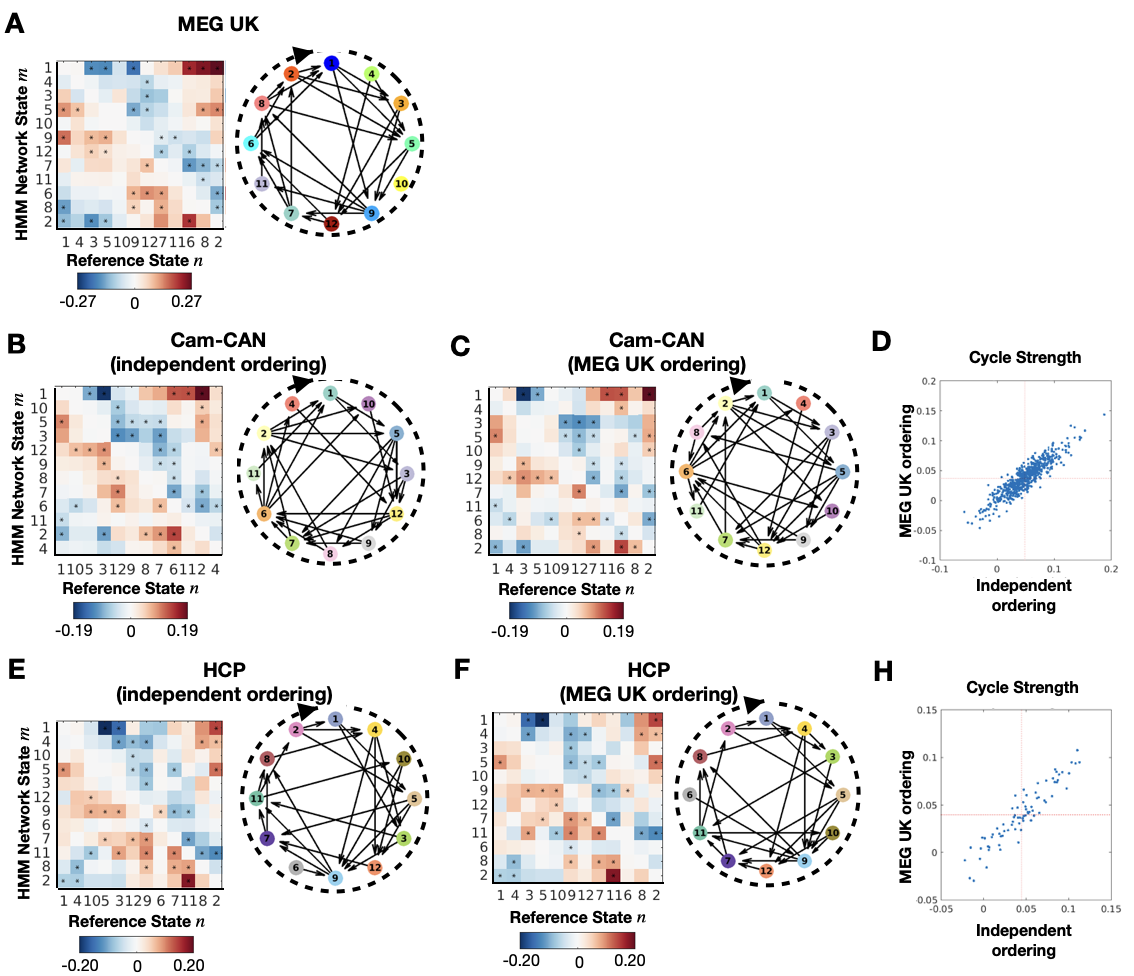

Figure S3. The overarching cyclical activation structure has equivalent ordering in all three datasets. A/B/E show the Fractional Occupancy (FO) Asymmetry (left) and cycle plot (right) where the ordering of states was optimised using the dataset under investigation. C/F show the same but with the MEG UK ordering. D/H show the quantitative agreement between an individual’s cycle strength using either ordering (dots show cycle strength for each subject; red dotted lines denote group means).

#### Supplement IV. Cyclical activation of functional brain networks does not appear when assuming a structure that is either Markovian or assumes a fixed time-lag.

When we ran TINDA on state time courses simulated from the transition probability matrix, only a hint of the observed FO asymmetry was present for all states. Moreover, this became only apparent after simulating 100 times as much data as exists in the observed data (Fig S3A-C). We then simulated state time courses from a non-Markovian model, by modelling the state transition probability at each point in time as a multinomial regression model in which the dependent variables were the states observed up to T timepoints prior to the current observation. This reproduced cyclical patterns that were still only a fraction of the strength of the patterns in the real data (one way ANOVA; p=6.8e-37); but which also show a linear relationship of increasing cycle strengths for longer permitted time lags (f-statistic 27.8; p=5e-4).

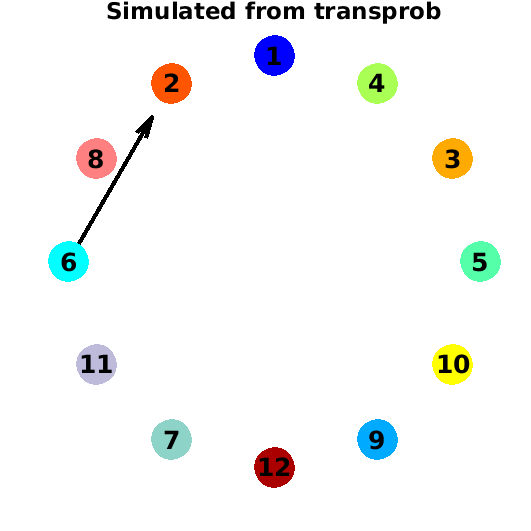

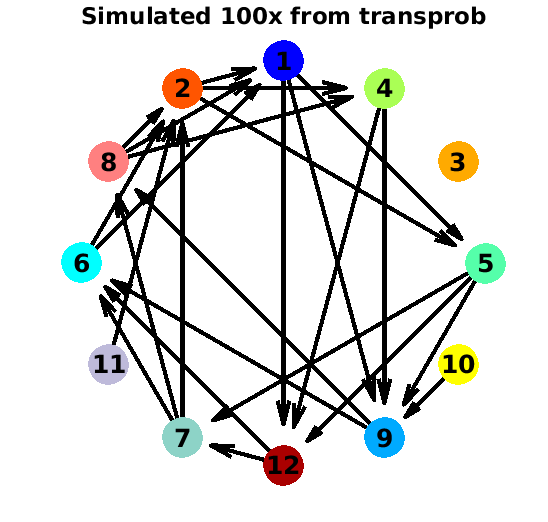

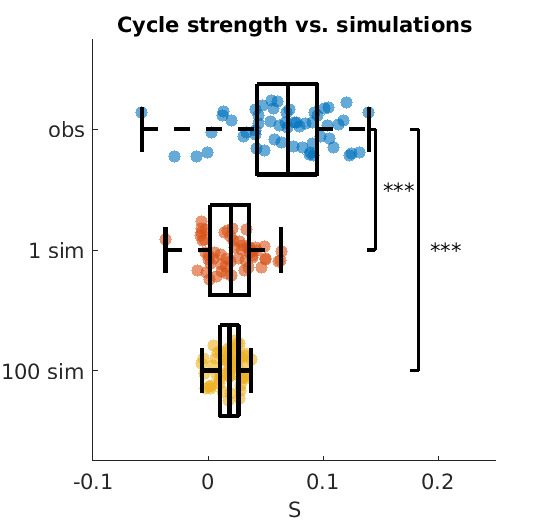

**C**

**B**

**A**

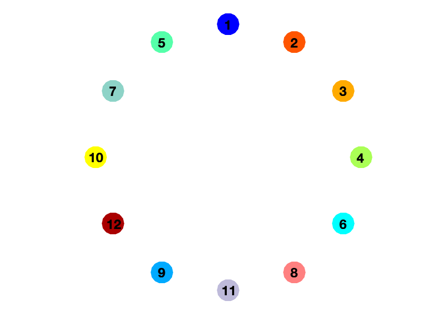

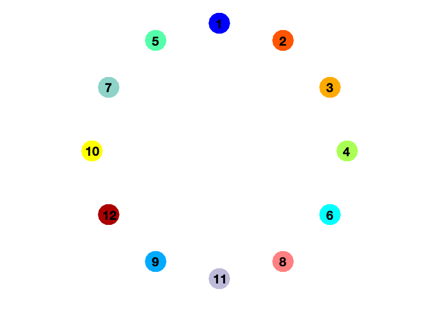

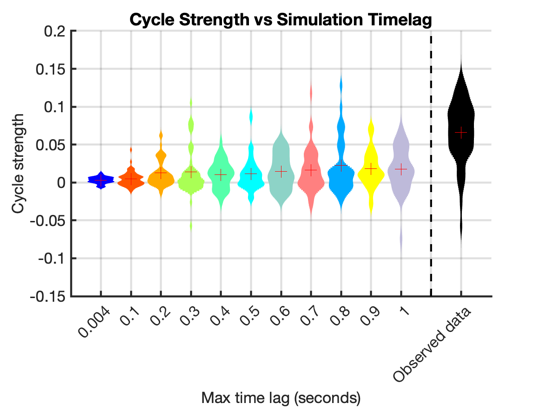

**D**

**E**

**F**

Figure S4. Cyclical activation of functional brain networks does not appear when assuming a structure that is either Markovian or assumes a fixed time-lag. A-B) Cycle plot when running TINDA on data simulated from the HMM transition probability, simulating the same (A) or 100 times (B) the amount of data in the observed data. C) Cycle strength is significantly lower in the simulated data. D-F) Similar to A-C, but for data simulated from a non-Markovian model.

#### Supplement V. Cyclical structure is strongest over timescales of seconds

We repeated the analysis in Figure 3 for the Cam-CAN, and HCP datasets and found consistent results. In summary, when we separate inter-state intervals (ISIs) into quintiles and apply TINDA to each quintile separately, we find that no cyclical structure is present at short interval times. Cycle strength is most strongly present at interval times of hundreds of milliseconds to multiple seconds.

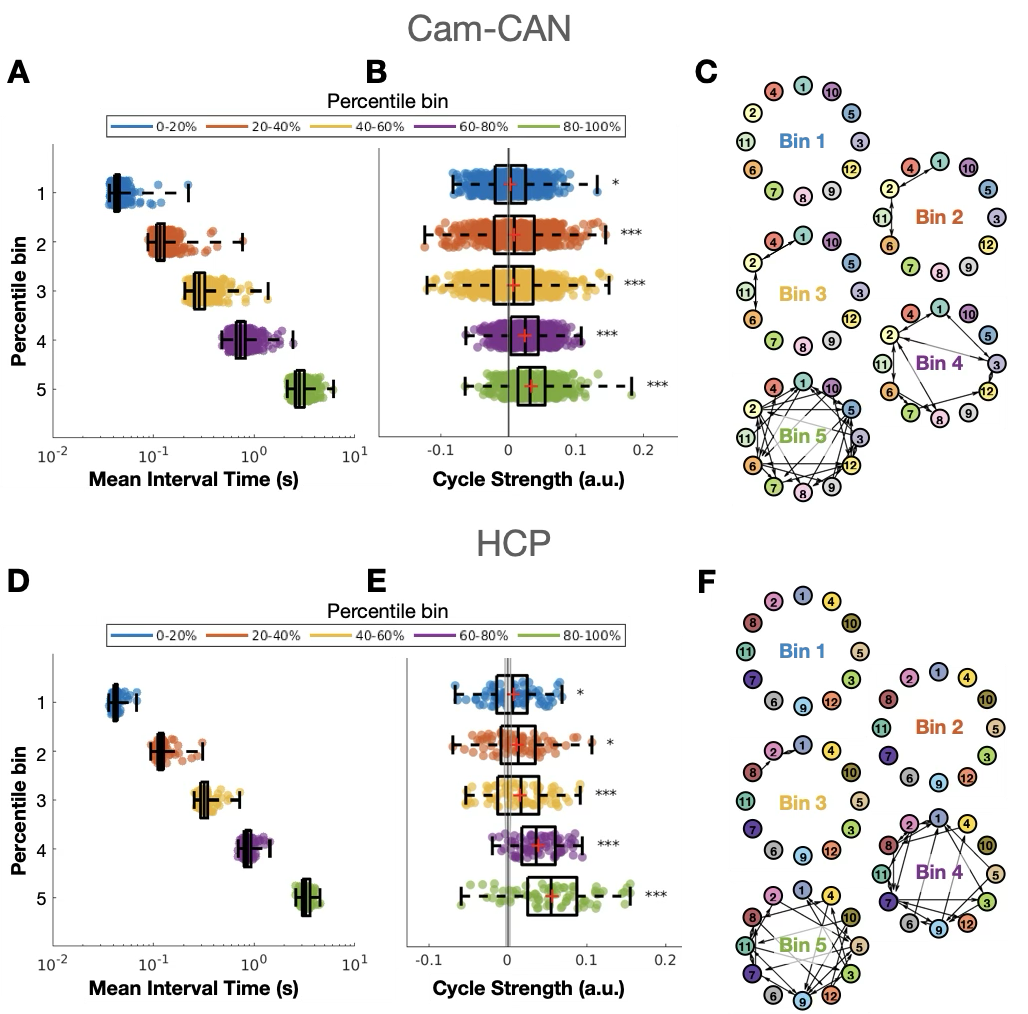

Figure S5. The observed cyclical organization of network state activations is driven by longer interval times. Replication of figure 3 for the Cam-CAN (A-C) and HCP (D-F) datasets. A/D) The mean duration of interval times within each percentile bin. B/E) The cycle strength resulting from running TINDA on each percentile bin in A. C/F) Graphs similar to Figure 2D for binned interval times with increasing duration from top to bottom. Circles are individual subjects; boxplots display the median, mean (+), 25^th^ and 75^th^ percentile, and whiskers indicate the minimal and maximal value. The line and error bar around zero cycle strength are the mean and standard deviation of the empirical permutation distribution. Significant cycle strength is denoted by asterisks: * p<0.05, ** p<0.01, *** p<0.001, and “n.s.” denotes not significant.

#### Supplement VI. Cyclical activation of functional brain networks are not artefacts of common physiological artifacts.

To make sure our results were not caused by other (rhythmic) physiological signals, we repeated the TINDA procedure in the MEG UK dataset on intervals spanning less than half the typical respiratory cycles (Figure S6A-B), and on intervals shorter than the individual’s typical heart rate (Figure S6C-D). While both procedures reduced the cycle strength substantially (with 23% and 65%, respectively) the cyclical pattern persisted in both and remained higher than expected by change (i.e., based on permutations). The reduction in cycle strength is in line with Figure 3 in the main manuscript, where we showed that the cyclical pattern is mostly present on longer time scales.

Moreover, to take these control analyses one step further and control for ocular activity, we also repeated TINDA on intervals of all durations but excluding segments within 500 ms of an identified ocular artefact (using the MNE algorithm *find_EOG_events*; proportion of data removed per subject: mean ± std = 22% ± 23%). We did this in the Cam-CAN dataset because here we had access to EOG data. Again, the cyclical pattern persisted (E), and cycle strength only reduced by 4.6%. On inspection, individual cycling characteristics remain highly correlated with their original values after removing ocular artefacts (F).
Together, these control analyses confirm that the cyclical patterns observed in the data are not a trivial consequence of other (rhythmic) physiological activity, and that they are likely a result of a genuine structure in neural network dynamics.

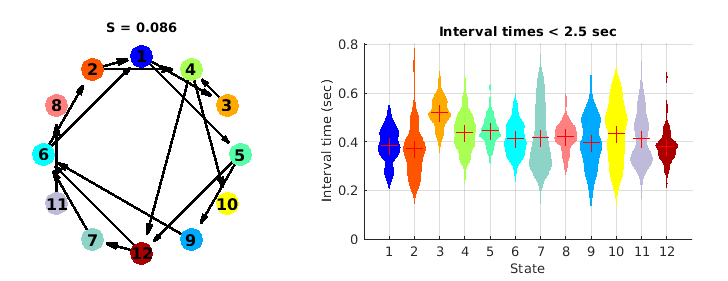

**B**

**A**

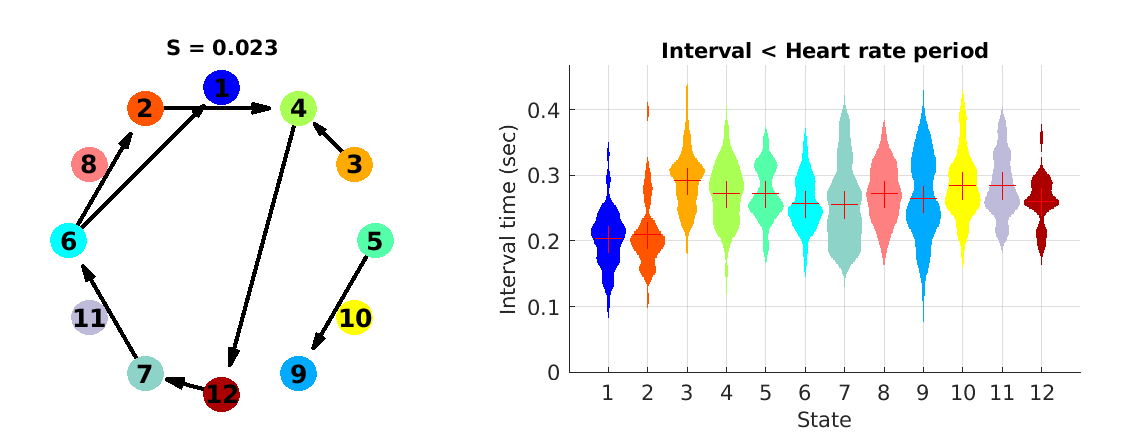

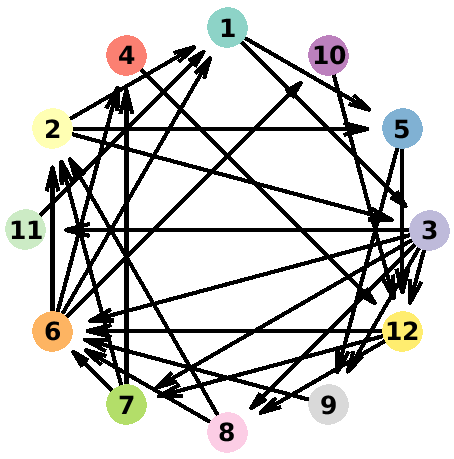

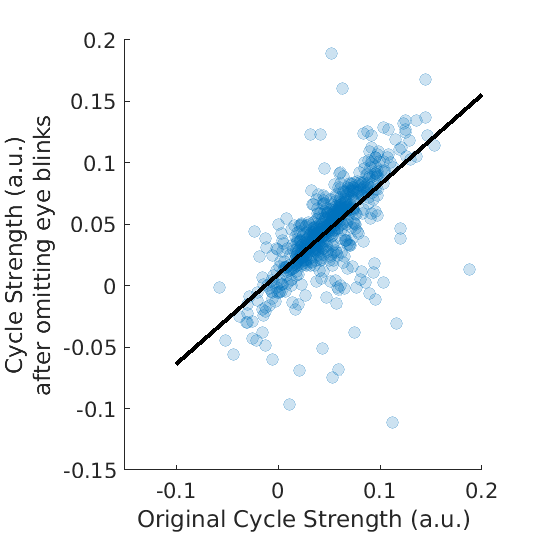

**D**

**C**

**E**

**F**

Figure S6. The Cyclical activation of functional brain networks is not a trivial consequence of cardiac or respiratory activity. The cyclical pattern resulting from applying TINDA to all intervals shorter than half a typical respiratory cycle. B) The distribution of mean interval durations over subjects used in A for each state. C) The cyclical pattern resulting from applying TINDA to all intervals shorter than an individual’s heart rate. D) The distribution of mean interval durations over subjects used in C for each state. E) The cyclical pattern resulting from applying TINDA to all intervals, excluding data within 500 ms of EOG artefacts. F) removing such data, which amounts to 20% of data, materially affects only a few subjects and does not affect the overall result.

#### Supplement VII. Spatial and spectral characterization of HMM state space along cyclical activation.

A	MEG UK

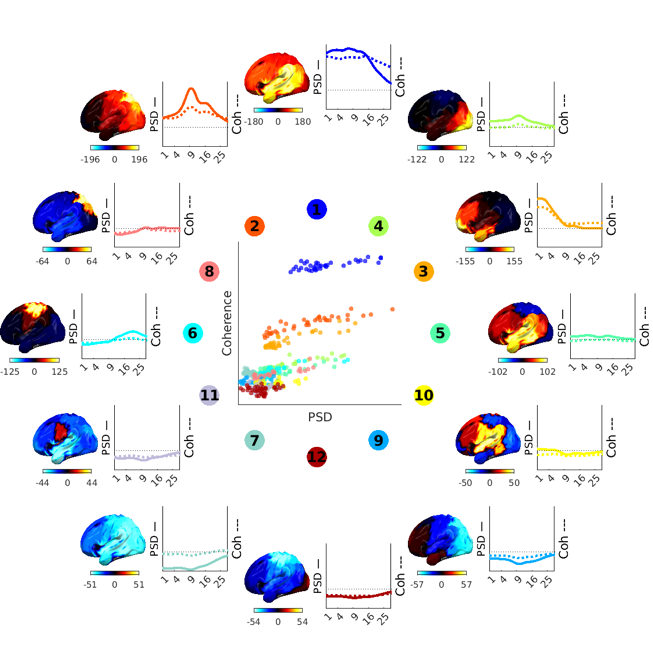

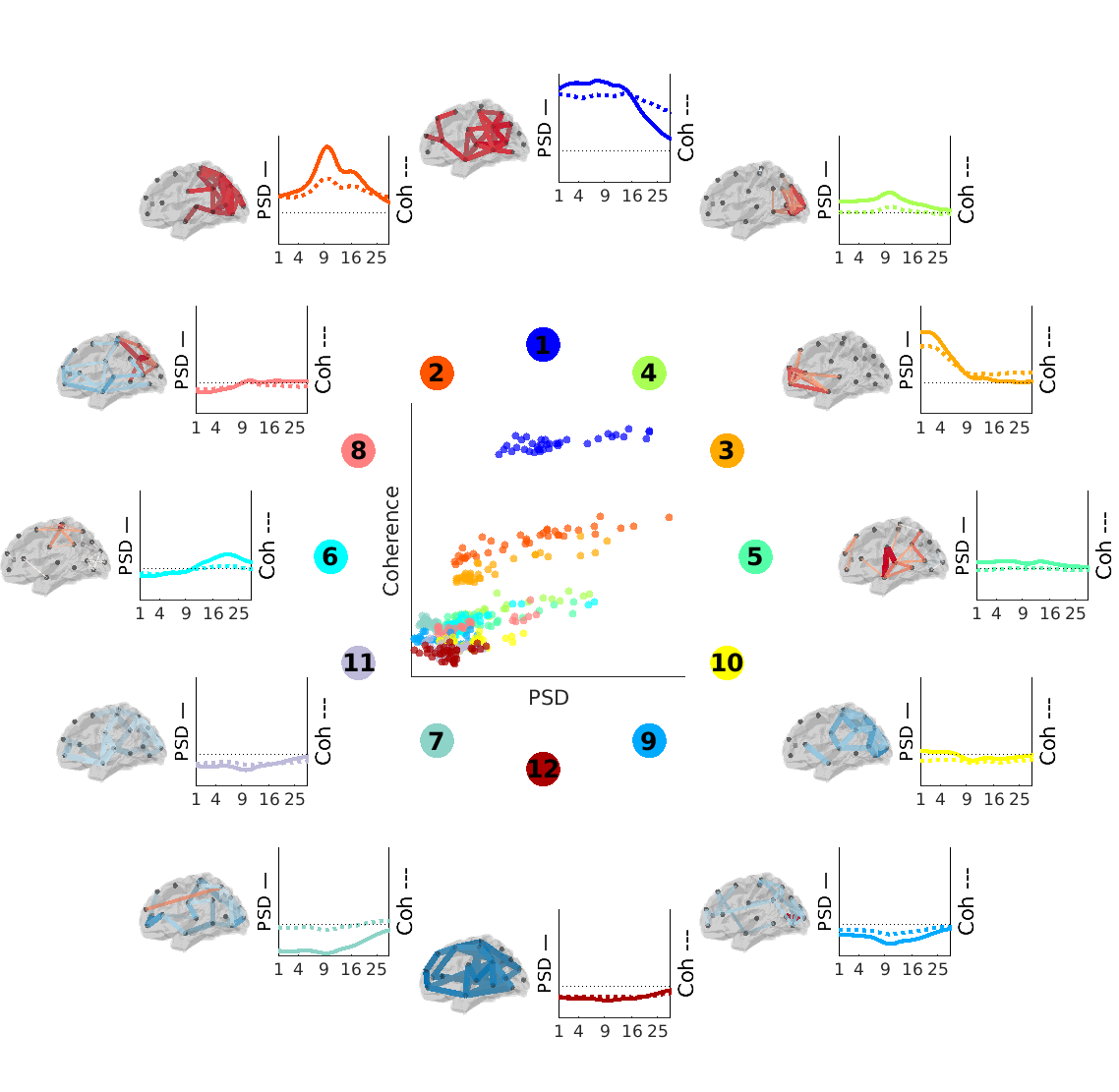

B	Cam-CAN

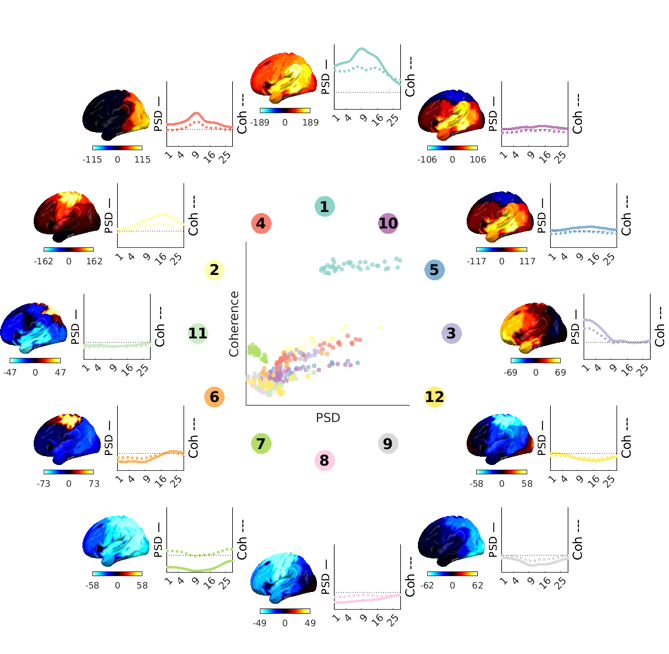

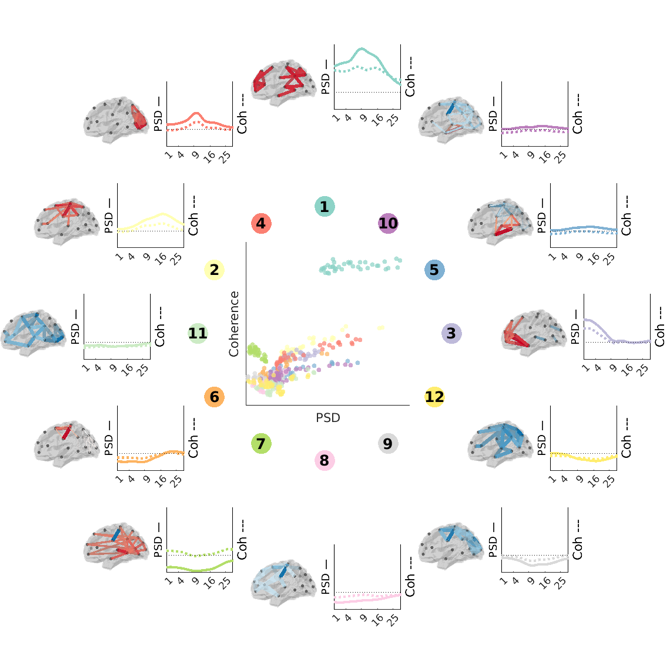

C	HCP

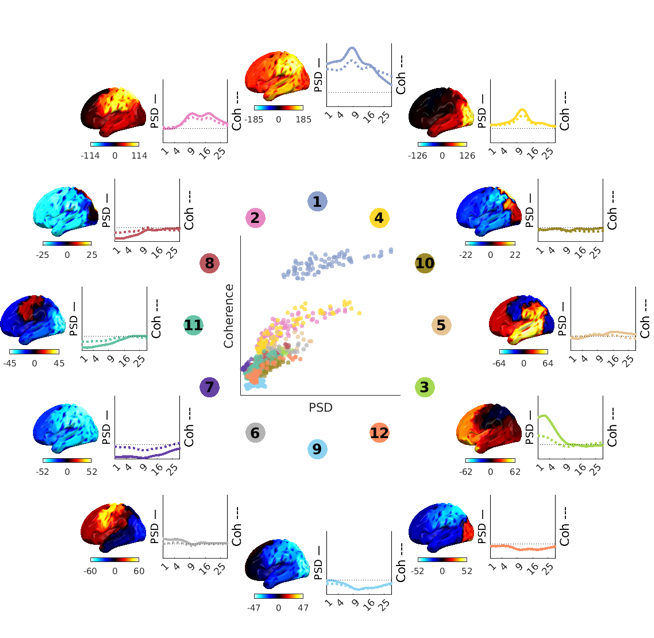

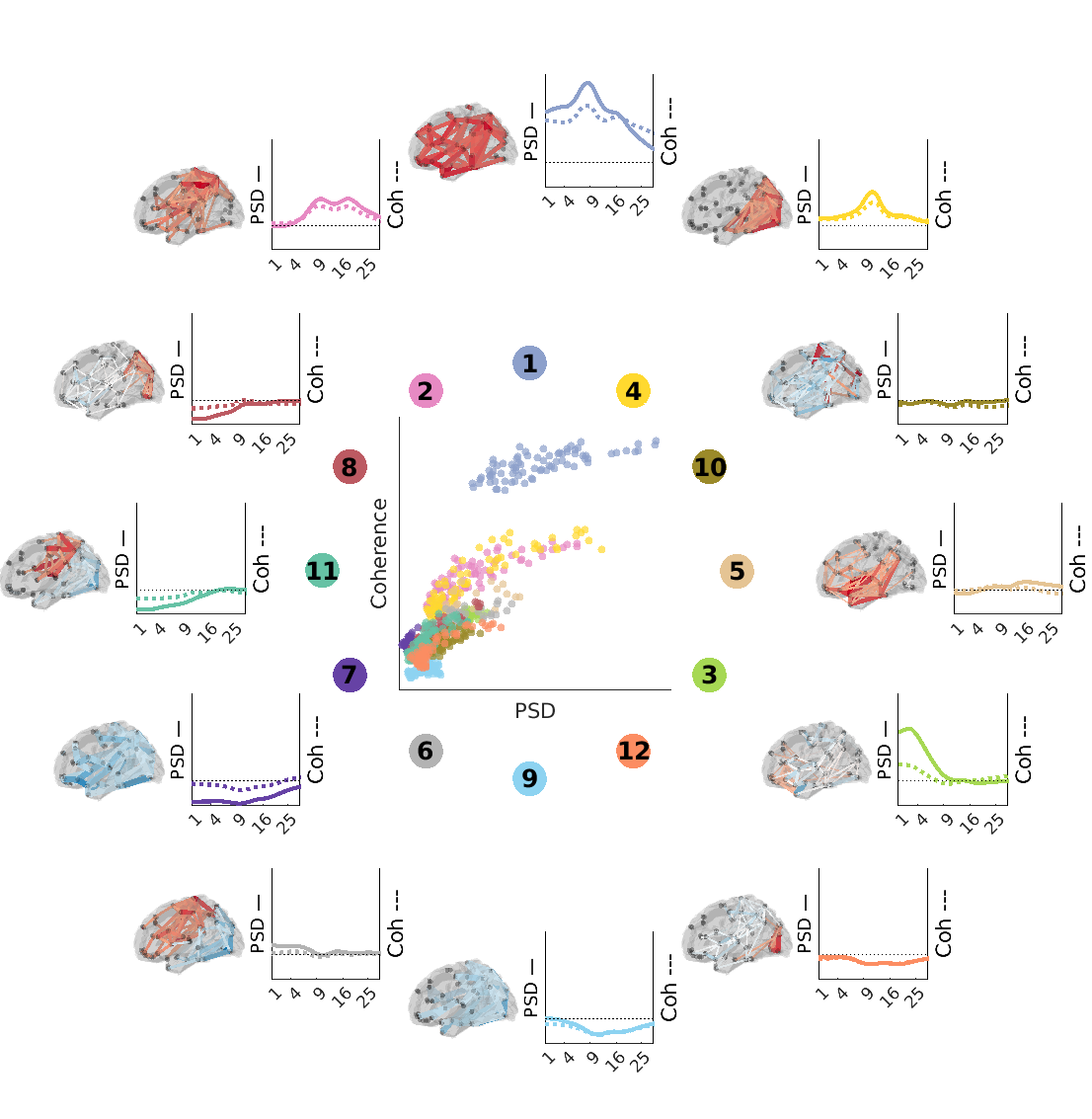

D	Wakeman-Henson

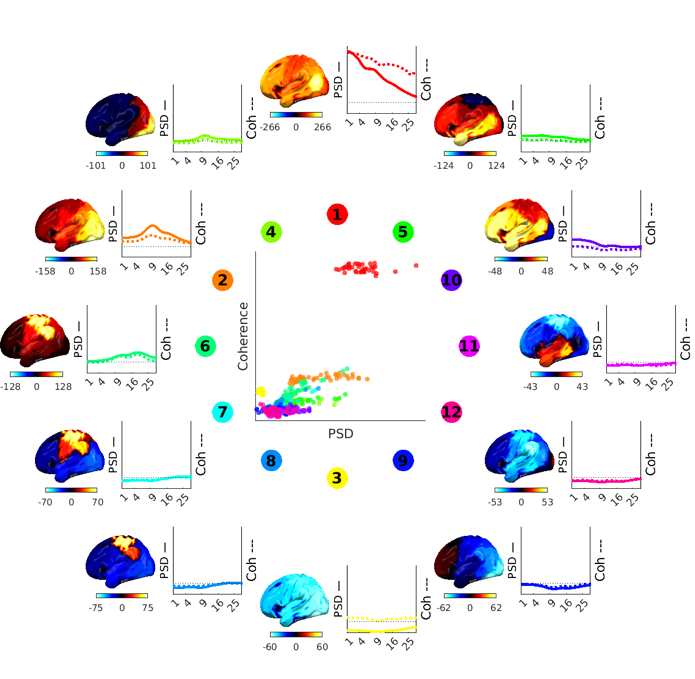

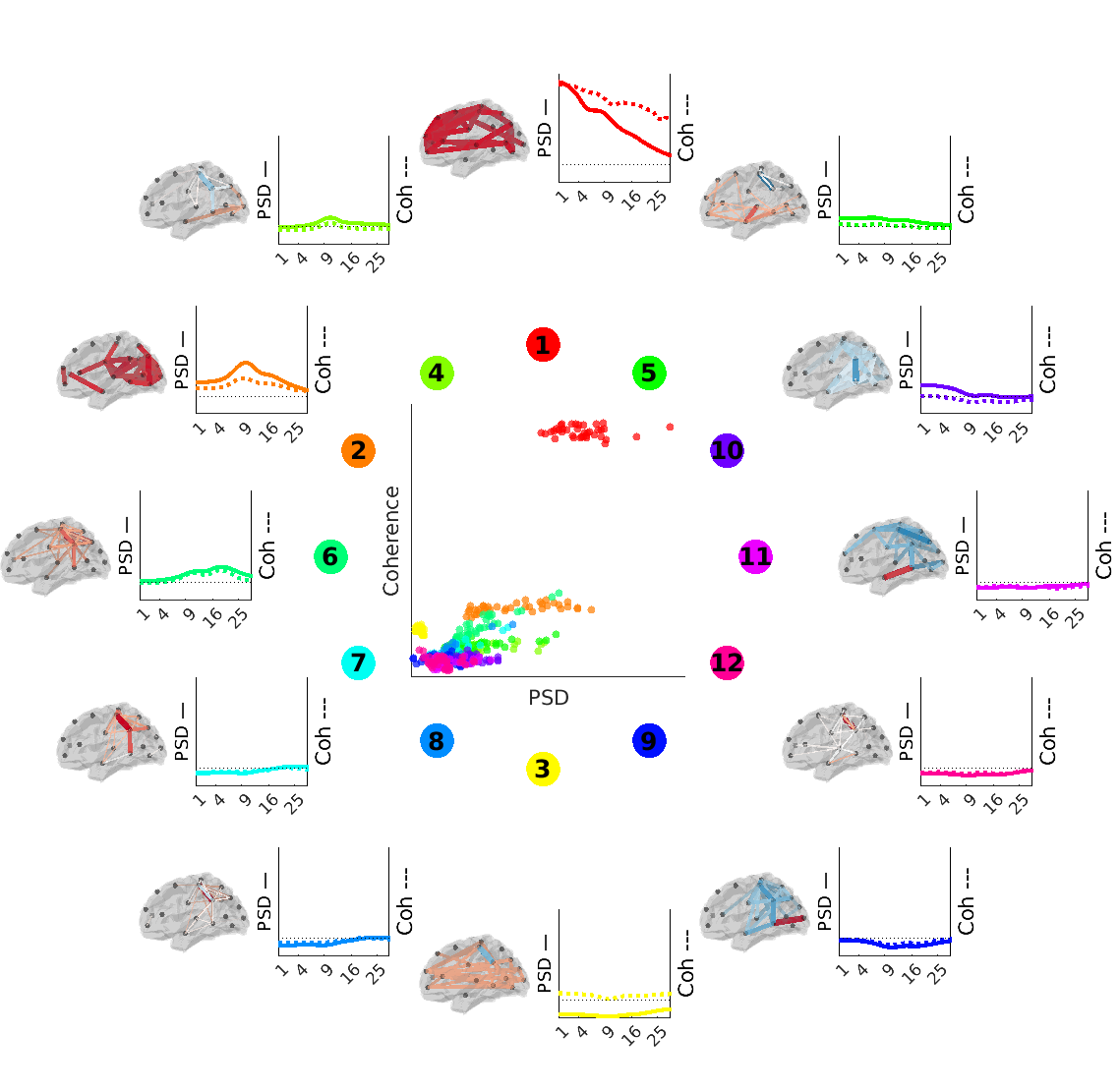

Figure S7. Spatial and spectral characterization of HMM state space along cyclical activation (as in Figure 4). Spatial maps of power (left) and coherence (right), for MEGUK (A), Cam-CAN (B), HCP (C), and Wakeman-Henson (D). Each brain map shows the power (coherence) increase (red) or decrease (blue) from the static power (coherence). To the right of each brain map is the spatial average PSD (solid line) and coherence (dotted line) as a function of frequency, relative to the static measures (horizontal dotted red line). The middle plot depicts the broadband average PSD versus coherence for each state (colour) and parcel (dot).

#### Supplement VIII. Characterizing individual cycle traversals

Inspired by the qualitative segmentation of the cycle into four “meta states” of distinct spatio-spectral characteristics (Figure 4B-C), we define a full cycle as being made up of a sequential activation of these four meta states, which we infer using a second level HMM run on the time courses of sliding window fractional occupancy computed on the network state time-courses obtained from the first level HMM described thus far (see Methods for details).

With this approach we sought to characterise what a ‘typical’ traversal of a cycle entailed in the MEG UK dataset. The group average mean cycle duration was 549 ms (Figure S8B), approximately 8.7 times longer than the mean duration of individual network states. Evaluating the network states visited on each individual cycle revealed these only comprised 5.4 unique state visits on average (Figure S8B). This suggests that a typical cycle traversal visits less than half of the K=12 available network states, but returns to several of those states at least once, i.e., through backward transitions within a meta state.

Plotting the phase shifts of each observed state transition further reinforces the view that cyclical dynamics emerge from stochastic and noisy short-term network state dynamics. If we consider each state transition on the cyclical clock face as a phase shift in the complex plane, we can build a histogram of these transitions as in Figure S8D. The mode of this plot is a state transition that is one step in the clockwise direction. However, a state transition two steps clockwise is as probable as a transition one step counterclockwise; thus, a net clockwise motion is achieved only with frequent steps back in the counterclockwise direction. Furthermore, jumps of four or more steps in either direction – that we would characterise as broadly ‘acyclic’ transitions – nonetheless account for 25% of all state transitions. In conclusion, the cyclic dynamics we have identified emerge from noisy short term state transitions.

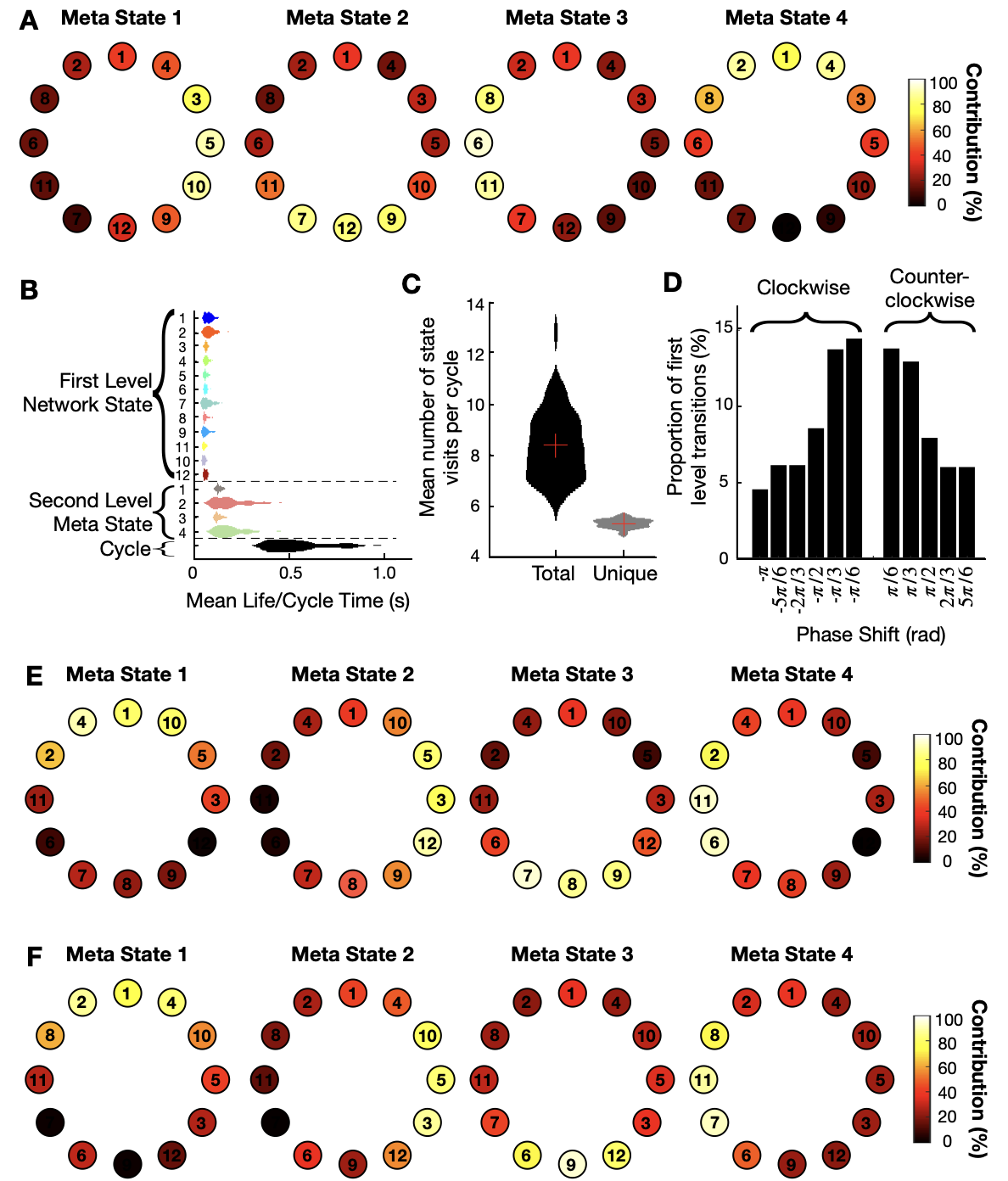

Figure S8: Characterizing individual cycle traversals. A) The contribution of each first-level HMM network state to the second level HMM meta states (Figure 4B-C) that are activated sequentially in one cycle traversal. B) Distribution of first and second level HMM state lifetimes and cycle times over subjects. C) Distribution of the mean number of states visited in each cycle over subjects. D) Histogram of individual state transitions represented as phase-shifts on the sequential clock face; clockwise transitions are represented as negative phase shifts, whereas anticlockwise transitions are represented as positive phase shifts. E-F) Similar to A, but for the Cam-CAN (E) and HCP (F) studies.

#### Supplement IX. Robustness of findings over parameters

In the main manuscript we have shown the presence of the cyclical structure and the ordering of states within it is reproducible across datasets. Here, we show the dependency of cycle summary metrics and their correlation with demographics on the number of network states fitted in the first level HMM. For this, we used the Cam-CAN dataset, because it has the most subjects (which improves the robustness of individual HMM fits), and because of the correlations with age shown in the main manuscript (Figure 5). We fitted 4 to 18 network states. Lastly, we test the dependency of the heritability of cycle metrics to various confounds, including, age, sex, and morphometric factors (Figure S10).
We investigated the dependence of cycle rate and cycle strength on the number of network states, as well as the correlation over subjects of the metric with the original presented in the main manuscript, and the correlation over subjects with age (Figure S9). Mean cycle strength was much lower for 4 states, but the strength at 5-18 states was stable and showed a slight downward trend with the number of states (Figure S9A). The correlation with the cycle rate of the original, 12-state model was >0.7 for 5-8 states, >0.8 for 9 states and higher, and highest for models with 9-14 states, and the correlation with age was remarkably stable for all models (Figure S9B/C). We observed similar trends in the cycle rate, albeit more noisy (Figure S9D-F). In particular, the absolute value of the cycle rate and the correlation with age are more variable between models where the difference in the number of states fitted was small. This is not surprising given the extra complexity of the second level HMM. In conclusion, the cycle metrics described in the main manuscript are robust to the number of network states fitted in the first level HMM, especially when 8 or more states are fitted.

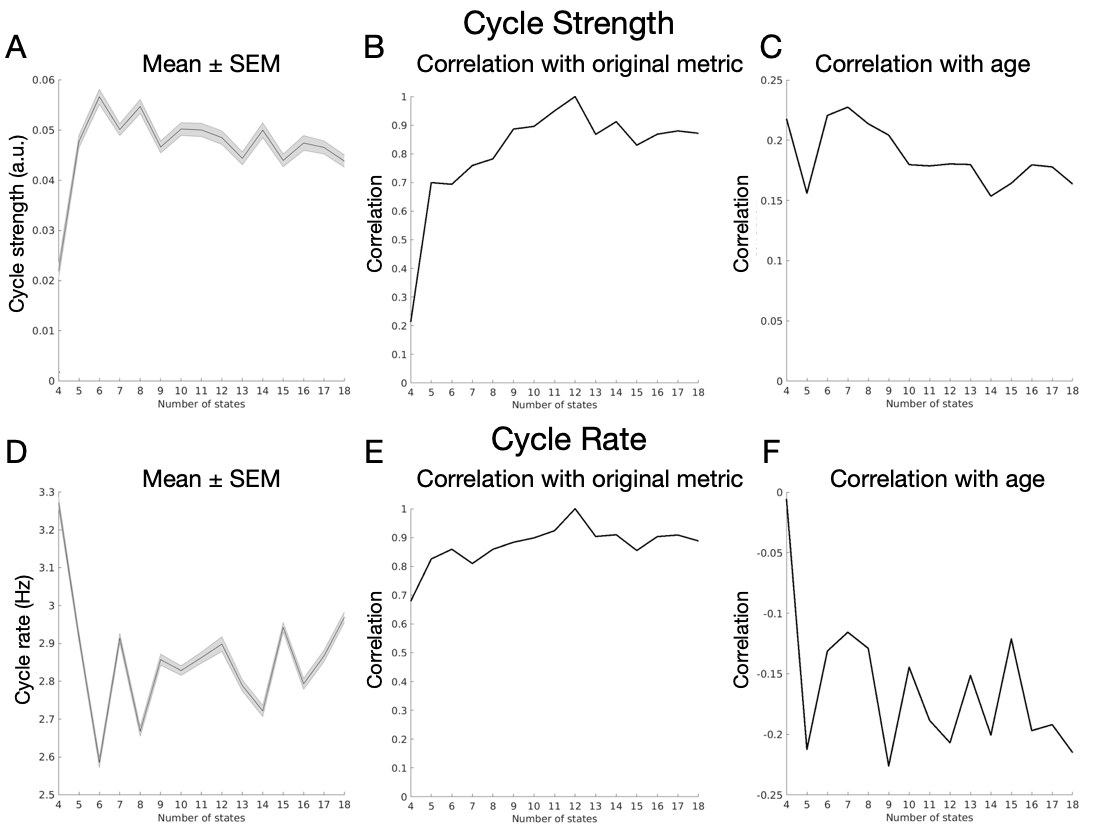

Figure S9. Cycle metrics are robust to the number of network states fitted in the first level HMM. A) The mean and mean standard error (SEM) of cycle strength as a function of the number of states fitted. B) The correlation over subjects with cycle strength in the original, 12 state model, and C) the correlation with age. D-F) Similar to A-C but for cycle rate.

The associations between cycle metrics and demographic variables are also robust to minor changes in the selected metrics. In the interests of transparency, we report that we tested a small range of metrics to characterise cyclical dynamics. As shown below, these tended to reflect largely overlapping information with either cycle rate or cycle strength (i.e. with the two metrics reported in the main manuscript).
As an alternative to cycle strength, we defined “circularity” as a metric of how geometrically circular a directed graph is, i.e., computing the clockwise distances of the significant FO asymmetries, and comparing these with permutations. Here, the assumption is that a “perfect geometric circle” has small clockwise distances (e.g., from 12 o’clock to 1 o’clock) rather than the opposite (e.g., from 12 o’clock to 4 o’clock, or 11 o’clock). We abandoned this metric because it was less well-founded in comparison with cycle strength.
Still circularity was highly correlated with Cycle strength; and mean / median measures of cycle time / rate were highly correlated / anticorrelated with the mean cycle rate (Figure S9B). The metrics cycle rate and cycle strength were chosen as they were the most Gaussian of these correlated measures (as reflected by the Kolmogorov-Smirnov test statistic in figure S9C below).

Finally, Table S9 shows the regression model predicting age and sex reproduced for different selections of these parameters. None of the results presented in the manuscript are materially changed by selection of alternative metrics for measuring cyclical dynamics.

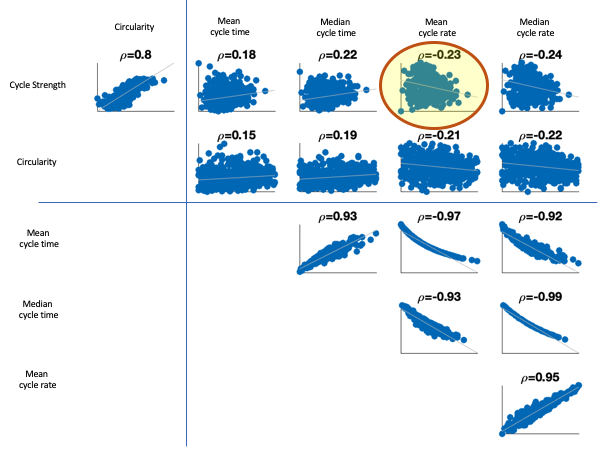

Figure S9B: alternative metrics to represent cycle rate and their mutual dependencies. Combination highlighted in yellow are the two metrics reported in the manuscript.

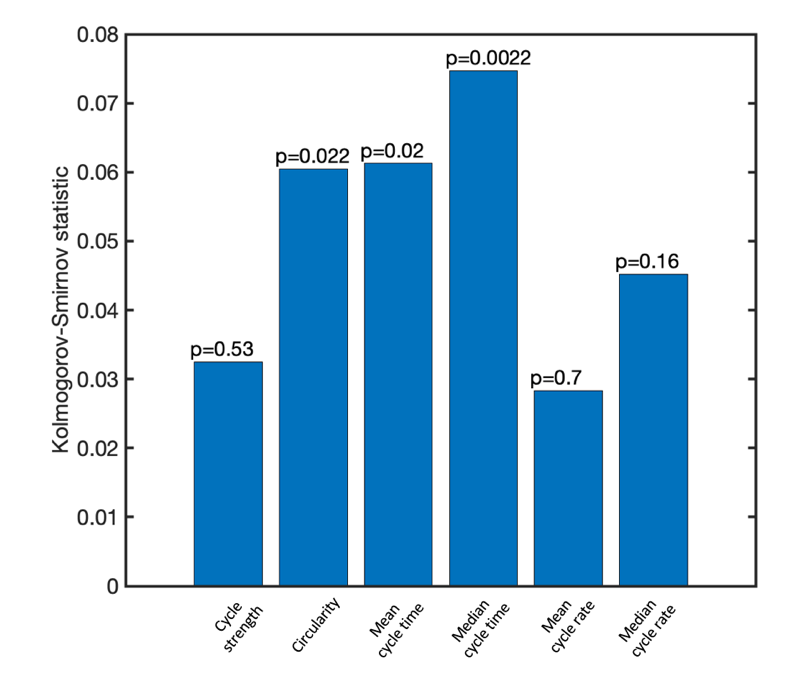

Figure S9C: metrics were selected (cycle strength and mean cycle rate) taking into account the relationships of Figure S9B and their Gaussianity, as measured by Kolmogorov-Smirnov test statistic.

Table S9A: robustness of multiple regression model predicting age effect to different metric selection. The regression model used to predict age allows us to substitute different metrics. The entry highlighted corresponds to the metrics reported in the main manuscript results; changes to metric selection do not materially affect the significance of any results reported.

| X2  X1 | Mean cycle period | Median cycle period | Mean cycle rate | Median cycle rate |
| --- | --- | --- | --- | --- |
| Cycle strength | B1=2.82 (p1=0.0002);  B2=1.65  (p2=0.028) | B1=2.35  (p1=0.0017)  B2=2.96 (p2=0.0001) | **B1=2.49**  **(p1=0.0010)**  **B2=-2.04 (p2=0.0070)** | B1=2.25  (p1=0.0026)  B2=-3.17 (p2=2.7e-5) |
| Circularity | B1=3.36  (p1=6.6e-6)  B2=1.78 (p2=0.0165) | B1=2.98  (p1=5.6e-5)  B2=3.01 (p2=4.9e-5) | B1=3.18  (p1=1.9e-5)  B2=-1.93 (p2=0.0091) | B1=2.9 (p1=2.4e-5)  B2=-3.16 (p2=1.9e-5) |

Table S9B: robustness of multiple regression model predicting sex effect to different metric selection. The regression model used to predict sex allows us to substitute different metrics. The entry highlighted corresponds to the metrics reported in the main manuscript results; changes to metric selection do not materially affect the significance of any results reported.

| X2  X1 | Mean cycle period | Median cycle period | Mean cycle rate | Median cycle rate |
| --- | --- | --- | --- | --- |
| Cycle strength | B1=-0.052 (p1=0.53);  B2=-0.27  (p2=0.0013) | B1=-0.042  (p1=0.62)  B2=-0.26 (p2=0.0028) | **B1=-0.052**  **(p1=0.54)**  **B2=0.24 (p2=0.0050)** | B1=-0.035  (p1=0.68)  B2=0.28 (p2=0.001) |
| Circularity | B1=0.002  (p1=0.98)  B2=-0.29 (p2=6.5e-4) | B1=0.0044  (p1=0.96)  B2=-0.27 (p2=0.0017) | B1=0.011  (p1=0.9)  B2=0.29 (p2=6.8e-4) | B1=0.017 (p1=0.84)  B2=0.303 (p2=3.8e-4) |

The heritability estimate described in the main text (Figure 5) was particularly strong for cycle rate (h^2^=0.73) despite the limited amount of twin data present in the dataset (13 monozygotic, 11 dizygotic pairs of twins). Since we also observed age and sex effects on cycle metrics, these could inflate the heritability affect. In addition, another potential confound is brain and/or cortical volume. To assess the effect of these confounds, and for complete transparency, we here show the estimated heritability effect controlling for a different amount of potential confounds (as in ^1^).

We first estimated the heritability of cycle strength and rate in HCP without controlling for any confounds. Then, we controlled for the following confounds: sex, age, age^2^, sex × age, sex × age^2^, (brain volume)^1/3^, (cortical volume)^1/3^. We did so in a stepwise fashion, such that heritability was estimated after correcting (i.e., regressing out) sex, then sex and age, then sex, age, and age^2^, etc. Figure S10 shows that the heritability estimates of cycle strength and rate were relatively stable and only reduced slightly when controlling for more confounds. Importantly, the conclusion drawn from the results were equivalent: cycle strength is not heritable, cycle rate is heritable.

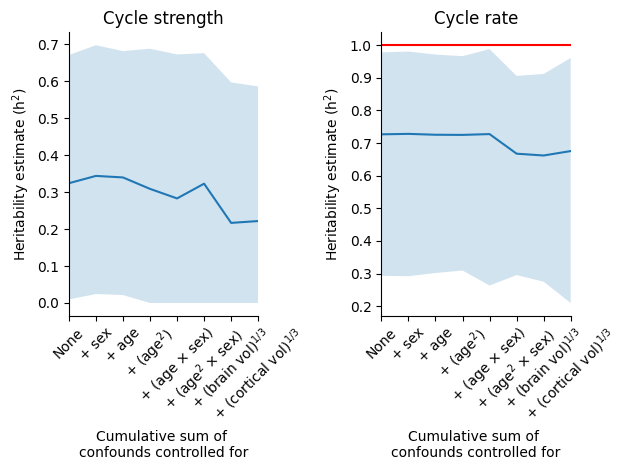

Figure S10. The robustness of cycle metric (A: cycle strength; B: cycle rate), heritability with regards to confounds controlled for. Confounds were controlled for (i.e., regressed out) in stepwise, cumulative fashion (x-axis), e.g., first only regressing out sex, then sex and age, then sex, age, and age^2^, etc. The solid blue line indicates the h^2^ estimate from the ACE model, with the coloured area indicating lower and upper bound, and red line significance at p<0.05).

#### Supplement X. Correlations of cycle metrics

In the main manuscript we showed significant correlations between cycle metrics with demographics and cognitive scores. Here, we replicated these analyses in other datasets where possible (i.e., where the dataset allowed it). To replicate the correlation analysis between cycle metrics and age and sex, originally shown in the Cam-CAN (N=612) dataset, we pooled the data across the remaining datasets (N=193). The age distribution in this combined dataset was centred on 30.3 years (mean; standard deviation=8.46; range 18-62), which is in stark contrast with the uniform age distribution in Cam-CAN (spanning 18-88 years). Despite this limitation, we reproduced the significant correlation between age and cycle strength (beta=1.51, t(190)=2.20, p=0.029) and cycle rate (beta=-1.76, t=-2.55, p=0.012), see Figure S11. The difference in cycle rate between males (N=86) and females (N=107) trended in the same direction as in Cam-CAN, but was not statistically significant (beta=0.14, t(190)=0.79, p=0.43), and neither was cycle strength (beta=-0.072, t(190)=-0.41, p=0.68). These analyses confirm the relationship between cycle metrics and age but call for further replications regarding the cycle rate difference between the sexes.
Other correlations between cycle metrics and demographics could not be directly replicated in other datasets. Neither of the other datasets contained cognitive scores that were directly comparable to Cam-CAN, and HCP was the only dataset with twin data, and thus the only dataset in which the heritability of cycle metrics could be tested.

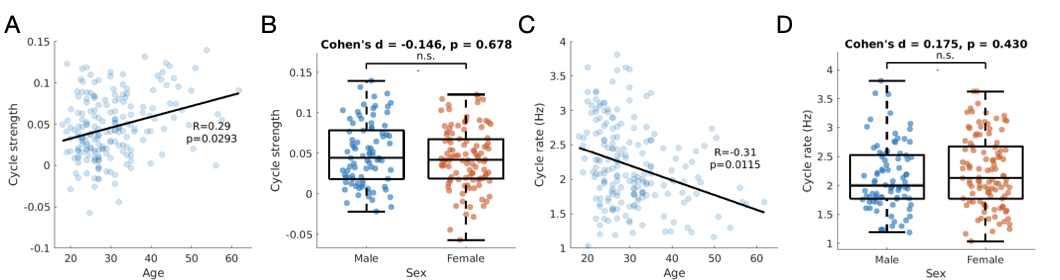
Figure S11. Correlations between cycle metrics and demographics in the pooled dataset (MEG UK, HCP, Replay, Wakeman-Henson), as in Figure 5. A) The correlation between cycle strength and age. B) Cycle strength for males and females. C-D) Similar to A-B but for cycle rate. Circles represent individual subjects. Boxplots display the median, 25^th^ and 75^th^ percentile, and whiskers indicate the minimal and maximal value.

Given that Cam-CAN has a high sample size and was designed to study age effects, we did post-hoc analysis on the Cam-CAN data for interpretational purposes. First, we found that cycle metrics (i.e., strength, and rate) are correlated (Fig S12A). We then wondered whether correlations between cycle strength and age can be explained by an overall increase in the size of Fractional Occupancy (FO) asymmetries. While this partly explains the results (Fig S8B), we also found that older people show more cyclical transitions compared to younger people. Next, we correlated mean switching rate of HMM states with age and found a significant correlation (Fig S12C). This indicates that the correlation between cycle rate and age might be caused by an overall increase in state switching. These strong links between cycle rate, cycle strength, and age warrant future research.

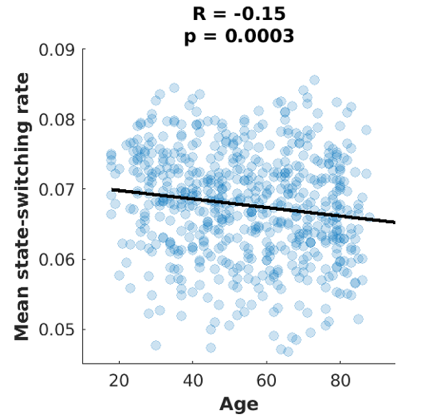

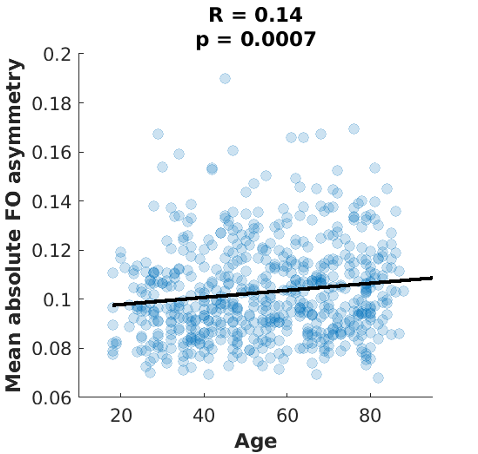

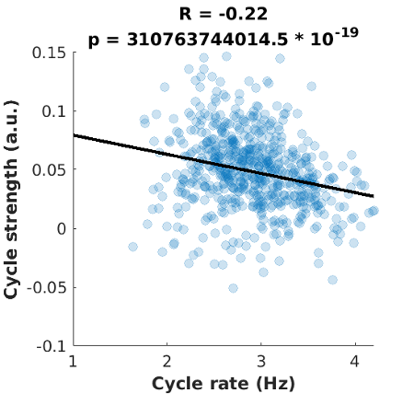

Figure S12. Correlations with and between cycle metrics. A) Cycle rate and cycle strength are correlated. Individuals who adhere to the cyclical structure more have faster cycle rates. Title refers to the statistic after regressing out age from both metrics. B) The mean absolute asymmetry between intervals in all states get larger with age. C) The mean switching rate decreases with age. Individual dots represent subjects in the Cam-CAN dataset.
Supplement XI. Individualised optimization of cycle order

In the main manuscript, we tested for the existence of a cyclical pattern on the group level. For example, the order of states was optimised based on the group level FO asymmetry. Cycle strength and cycle rate then relate to the group-level model of the cycle. This ambiguates the underlying causes for variability in cycle summary metrics. In particular, a low cycle strength could mean two things: 1) the optimal sequence order of the subject’s cycle is the same as the optimal order on the group level, but the subject’s brain network dynamics are not adhering to this cycle strongly (i.e., their dynamics are more stochastic), or 2) the optimal sequence order of the subject’s cycle is differs from the optimal order on the group level. If this is the case, the subject could adhere to its personal cycle order strongly or loosely; both would result in low cycle strength. Moreover, if a participant has strongly cyclical dynamics, but in a different order than the group level, the interpretability of cycle rate (i.e., according to the group-level cycle order) is in question.
To disambiguate these options, we did a post-hoc analysis in which we optimised the cycle order separately for each participant. We computed the cycle phase difference between each participant’s optimal cycle order and the optimal order estimated on the group level. We also computed the cycle strength and rate using the individual’s optimal cycle order.
We found both above options contribute to a low cycle strength in the main manuscript: participants with low cycle strength differed more in their optimal cycle order (R=-0.24, p=2.1 * 10^-9^; Figure S13A), but they still had low cycle strength using their optimal cycle order (R=0.67, p=6.7 * 10^-82^; Figure S13B). Importantly, the deviation from the group cycle order had little effect on the estimate of cycle rate: the cycle rate estimated using the group and individual order were strongly correlated (R=0.86, p=8.5 * 10^-184^; Figure S13C).
We also replicated the analysis from Figure S12A (i.e., correlation between cycle strength and rate) using the optimal individual order and found a strong negative correlation (R=-0.64, p=2.2 * 10^-72^; Figure S12D)

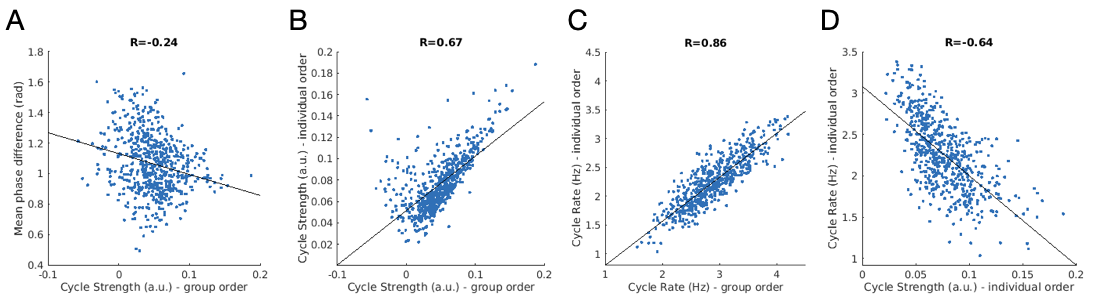

Figure S13. Correlations of summary metrics with group and personal cycle order optimization. A) The mean phase difference of state position between the individually optimised cycle order and the group order, as a function of cycle strength. B) The correlation of cycle strength from the individually optimised vs. group optimised cycle order. C) Like B but for cycle rate. D) as in Figure S12A but for the metrics estimated using the individually optimised cycle order.

#### Supplement XII. Correspondence of HMM states with fMRI resting state networks While resting state networks (RSNs) have been shown in M/EEG before, their correspondence to well-known RSNs, specifically the DMN and DAN from the fMRI literature, are here supported with quantitative comparison of the MEG HMM states to their fMRI equivalents. Given the different timescales of the fMRI states and MEG states, a strictly one-to-one correspondence should not be expected, with evidence rather that slower fMRI states comprise multiple faster MEG states^2^. We computed the dice coefficient^3^ between the MEG HMM state connectivity and the fMRI connectivity in the commonly used Yeo-7 DMN and DAN networks^4^. In their original publication, voxels were clustered into networks based on their common connectivity profile, where connectivity was estimated as the Pearson correlation between each voxel’s time course and those of regions of interest (ROI). We therefore mapped the connectivity profile for the Yeo-7 DMN and DAN into the same set of ROIs used in the MEG analysis, obtaining two 38x38 fMRI connectivity matrices representing the DMN and DAN connectivity profile. For the MEG HMM states, we defined connectivity by the wideband coherence for each state, leading to twelve 38x38 MEG connectivity matrices. Both MEG and fMRI connectivity matrices were averaged over subjects, and thresholded at the 90^th^ percentile. The dice coefficient between the thresholded matrices were computed and visualised below. Importantly, this confirms previous qualitative relationships between MEG and fMRI networks, namely the Default Mode Network corresponding most strongly to the HMM states 1 and 3 (top right phases of the cycle), and the dorsal attention network (DAN) corresponding to low power states 7, 9 and 11 (bottom left phases of the cycle).

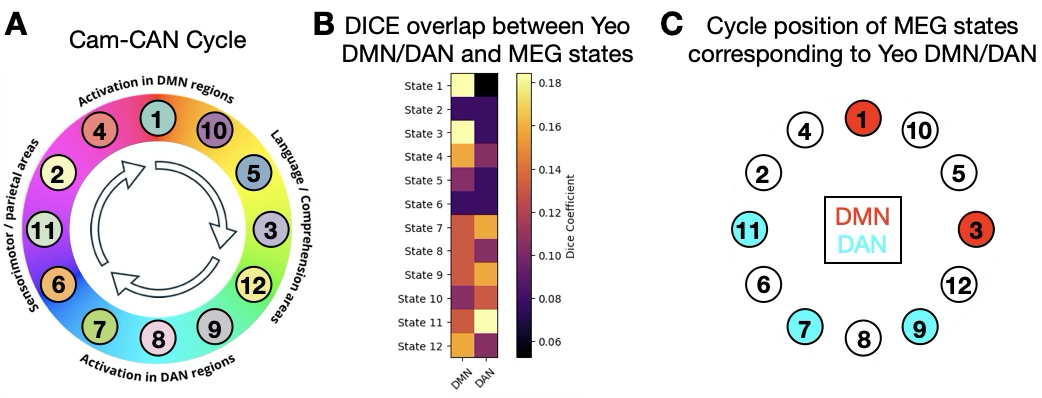

#### FigureS14. Dice scores between the Yeo-7 fMRI default mode (DMN) and dorsal attention networks (DAN) and the MEG HMM states. A) the conceptual model of Figure 4C, reproduced for the Cam-CAN dataset. B) DICE scores between Cam-CAN fMRI connectivity, and MEG connectivity. Focusing on the DMN and DAN networks, we see the fMRI DMN network connectivity has strongest correspondence with MEG states 1 and 3, corresponding to the top right quadrant of the cycle (C). Conversely, we see the fMRI DAN network connectivity has strongest correspondence with MEG states 7, 9 and 11, corresponding to the lower quadrant of the cycle. This is broadly consistent with the conceptual model (S14A and Figure 4C) linking the DMN and DAN as occupying opposing phases of the characterised cycle space.

### Supplement XIII. Correlations between cycle metrics and cognitive scores

In figure 5 of main manuscript, we established a relationship between cycle metrics (cycle strength, and rate), and 13 cognitive scores as present in the Cam-CAN dataset. Figure 5G showed the canonical weights for the second (significant) pair of canonical variates. Below, we show the canonical weights for the first pair of canonical variates (Figure S15A).
For interpretability and transparency, here we also report Pearson correlations between individual metrics post-hoc (Figure S15B). This shows that cycle strength has the strongest, negative correlation with verbal fluency, and cycle rate has negative correlations with fluid intelligence, motor speed (variance), and “spot the word”.

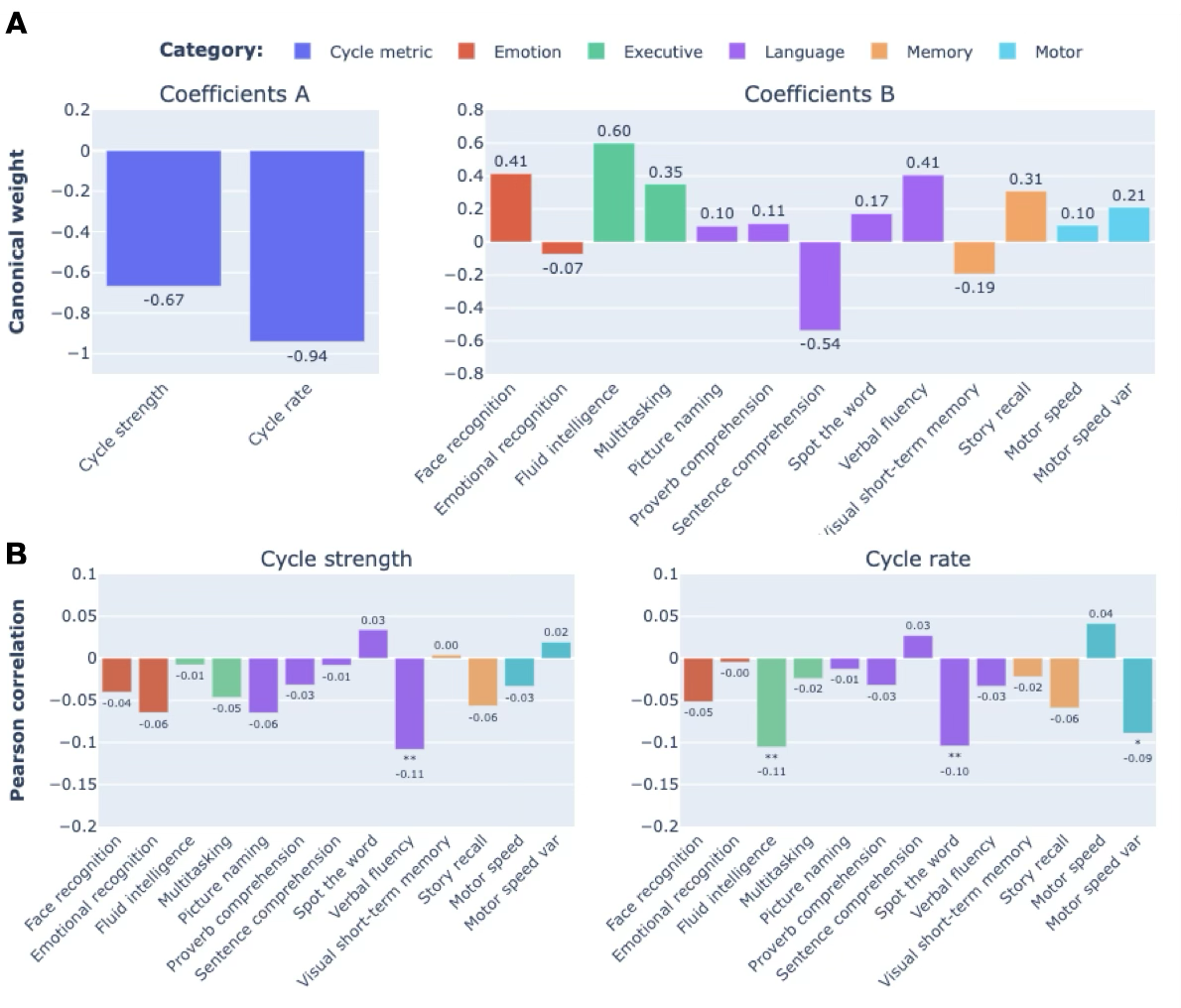

Figure S15. A) Canonical weights from the first (non-significant) pair of canonical variates; cycle metrics (left) and cognitive scores (right). B) Post-hoc Pearson correlations between cognitive scores and cycle strength (left) or cycle rate (right). Asterisks denote significant, uncorrected correlations (* p<0.05, ** p<0.01).
